# A computational framework linking synaptic adaptation to circuit behaviors in the early visual system

**DOI:** 10.1101/2022.08.27.505287

**Authors:** Liuyuan He, Yutao He, Kehuan Lun, Lei Ma, Kai Du, Tiejun Huang

## Abstract

Retina ribbon synapses are the first synapses in the visual system. Unlike the conventional synapses in the central nervous system triggered by action potentials, ribbon synapses are uniquely driven by *graded* membrane potentials and are thought to transfer early sensory information faithfully. However, how ribbon synapses compress the visual signals and contribute to visual adaptation in retina circuits is less understood. To this end, we introduce a physiologically constrained module for the ribbon synapse, termed Ribbon Adaptive Block (RAB), and an extended “hierarchical Linear-Nonlinear-Synapse” (hLNS) framework for the retina circuit. Our models can elegantly reproduce a wide range of experimental recordings on synaptic and circuit-level adaptive behaviors across different cell types and species. In particular, it shows strong robustness to unseen stimulus protocols. Intriguingly, when using the hLNS framework to fit intra-cellular recordings from the retina circuit under stimuli similar to natural conditions, we revealed rich and diverse adaptive time constants of ribbon synapses. Furthermore, we predicted a frequency-sensitive gain-control strategy for the synapse between the photoreceptor and the CX bipolar cell, which differ from the classic contrast-based strategy in retina circuits. Overall, our framework provides a powerful analytical tool for exploring synaptic adaptation mechanisms in early sensory coding.

## Introduction

Imagine walking through a dense forest now: the sun shines deep into the forest, the ground is full of spots of sunlight, squirrels and hares are jumping in this light and dark environment, and there may be snakes lurking in the grass. Our sensory systems are constantly bombarded with rich stimuli from this vivid natural environment. To adapt to the quickly changing environment, the visual sensory system needs to compress the enormous inputs from the surrounding world into meaningful electrical signals and project them to the higher visual center. This process involves a critical phenomenon in our visual perception, “visual adaptation” (1, 2). In the early visual system, adaptation strengthens weak luminance against noise and prevents response saturation to intense luminance. Specifically, the retina uses adaptive strategies to transform large-contrast or high-frequency light stimuli into neural signals(3-5). Visual adaptation in the retina involves intrinsic mechanisms at the cellular and circuit levels, such as phototransduction, synaptic transmissions, spike generations, and the inhibition of interneurons(2, 6). Among these mechanisms, signal transmissions by non-spiking chemical synapses, particularly the ribbon synapse, significantly contribute to adaptation(7). Ribbon synapses exclusively exist in early sensory systems and the pineal gland for graded signal encoding(8, 9). At the system level, retinal ribbon synapses are involved in various adaptive functions, including the extension of luminance range(10), high-frequency filtering(11), and the accurate encoding of light changes(12).

Yet it has taken over a half-century to gradually reveal adaptation-related sub-cellular mechanisms since the discovery of synaptic ribbons in the 1960s(13, 14). Specifically, the ribbon synapse’s Rapidly Releasable Pools (RRP) can rapidly respond to the increase in membrane potential as vesicles in the RRP are located close to the Active Zone(12, 15). Notably, the depression of ribbon synapses, a phenomenon strongly related to cell-level adaptation, is mainly induced by the fast depletion of vesicles in the RRP(12, 16, 17) and only moderately regulated by reducing inward calcium currents(18, 19), distinct from depression mechanisms in central synapses(20, 21). However, it remains unclear how cellular mechanisms contribute to adaptive behavior at the circuit level due to the technical challenges of simultaneously monitoring subcellular activities and circuit behaviors(7).

Theoretical frameworks have been demonstrated to play essential roles in bridging the gap between synaptic activity and circuit-level adaptation. The prevailing computational models are the Linear-Nonlinear (LN) model and its extended circuits models(22-26). These abstract models have been successfully applied to large-scale neural circuits and general information processing. However, the LN-based abstract models rely on data fitting and are not biophysically constrained, which may lead to their failure to capture adaptive behaviors when unseen stimuli are presented. Alternatively, recent works use biophysical models to account for the cellular basis of ribbon synapses in neural circuits(27-30). These detailed biophysical models exhibit improved biophysical interpretability, elegantly depicting the cascade of vesicle movements in presynaptic terminals. However, the design of these biophysical models requires precise knowledge beyond the reach of experimental techniques, such as vesicle resupply processes from the cytoplasm to the RRP(31-33). In addition, it’s infeasible to perform theoretical analysis on these models due to their complicated kinetics, severely preventing them from being widely used as an analytical tool in adaptation research.

Here, we developed a physiology-constrained module for the ribbon synapse, termed “Ribbon Adaptive Block (RAB),” and an extended circuit model, the “hierarchical Linear-Nonlinear-Synapse” (hLNS) model, to characterize the subcellular and circuit levels’ adaptive properties. Specifically, inspired by the classic Hodgkin-Huxley formalism(34), the RAB module follows the 2-state transition kinetics, which can theoretically be derived from the biophysical processes of vesicle release and resupply in ribbon synapses, ensuring a high degree of interpretability of the model. Notably, as a powerful theoretical tool, the RAB module analytically links vesicle movements to the adaptive properties, such as the adaptive time constant and the gain-control. Furthermore, our models successfully captured a variety of experimental recordings across different cell types and species, including the vesicle release on salamander cone synapses(12), the postsynaptic response on rat rod bipolar cell synapses(16), and the light-stimulated responses on mouse cone bipolar cell synapses(35). In particular, our model shows strong robustness to unseen stimuli, suggesting that it does capture the underlying mechanism of ribbon synapses, albeit simplistically compared to previous detailed models.

Next, with the hLNS model, we uncovered enriched adaptive time constants of ribbon synapses in the Photoreceptor – Bipolar Cell (BC) – Ganglion Cell (GC) circuit under stimuli closer to the natural conditions(35). Further, our theoretical analytical tool enables us to understand gain-controls under the stimuli with varied temporal frequencies and quickly changed contrasts, which is difficult to quantify experimentally. Intriguingly, we predict a frequency-sensitive gain-control strategy for the synapse between the photoreceptor – *C*_*X*_ bipolar cell, an atypical ON-type BC with a giant dendrite field (36), and few non-invaginating contacts to cones (37). Our finds are significantly complementary to the conventional contrast-based gain-control(38).

In conclusion, the RAB module, our theory-based and physiologically constrained model for ribbon synapses, successfully grabs synaptic adaptations at the circuit level and successfully uncovers adaptation strategies at several natural conditions, fulfilling the blank of experiments. We show RAB’s extensibility by using it as a simple yet powerful building unit in various circuit models containing ribbon synapses and generating adaptive behaviors consistent with empirical findings. Thus, with its simplicity that facilitates further analyses and the capability of capturing synaptic adaptation, RAB can be served as a lightweight analytical tool in relating the subcellular-level adaptive behaviors to a broad range of circuit-level adaptation.

## Results

As a particular type of neuronal synapses, ribbon synapses share some features with conventional synapses while differing mechanisms are underlying neurotransmitters’ release. Notably, at the step of transmitter release, we can observe a fast-to-slow switch of the release rate in both central and ribbon synapses, which is an essential adaptive behavior of chemical synapses(39, 40). In central synapses, two separate releasable pools or a single releasable pool with two distinct release components contribute to the adaptation(41, 42). In contrast, recent studies support the idea that the vesicle release with a unique mechanism in only one releasable pool, the RRP, can produce adaptive responses in ribbon synapses(12, 16, 29). It is due to a specific structure (the synaptic ribbon) in the ribbon synapse, which stores vesicles immediately behind the RRP, supporting most vesicle transmissions(15, 43-45). However, the vesicle depletion is still significant and produces intrinsic adaptive behaviors of ribbon synapses. Therefore, our synaptic adaptation module aims to use a unique response mechanism to model adaptive behaviors in an isolated ribbon synapse and a ribbon synapse inside a retinal circuit.

### The Ribbon Adaptive Block Module

We first utilize a two-state transition module obeying first-order kinetics to capture the release process of neurotransmitters in ribbon synapses so that the module endows the intrinsic synaptic adaptive properties. Specifically, we use two states to describe vesicle sites in the RRP. We suppose a vesicle’s release and resupply processes are instant so that at one moment, a vesicle site is occupied with vesicles or not. Thus, the Ribbon Adaptive Block(RAB) has the following diagram (Fig.2A upper middle):

**Figure 1.**
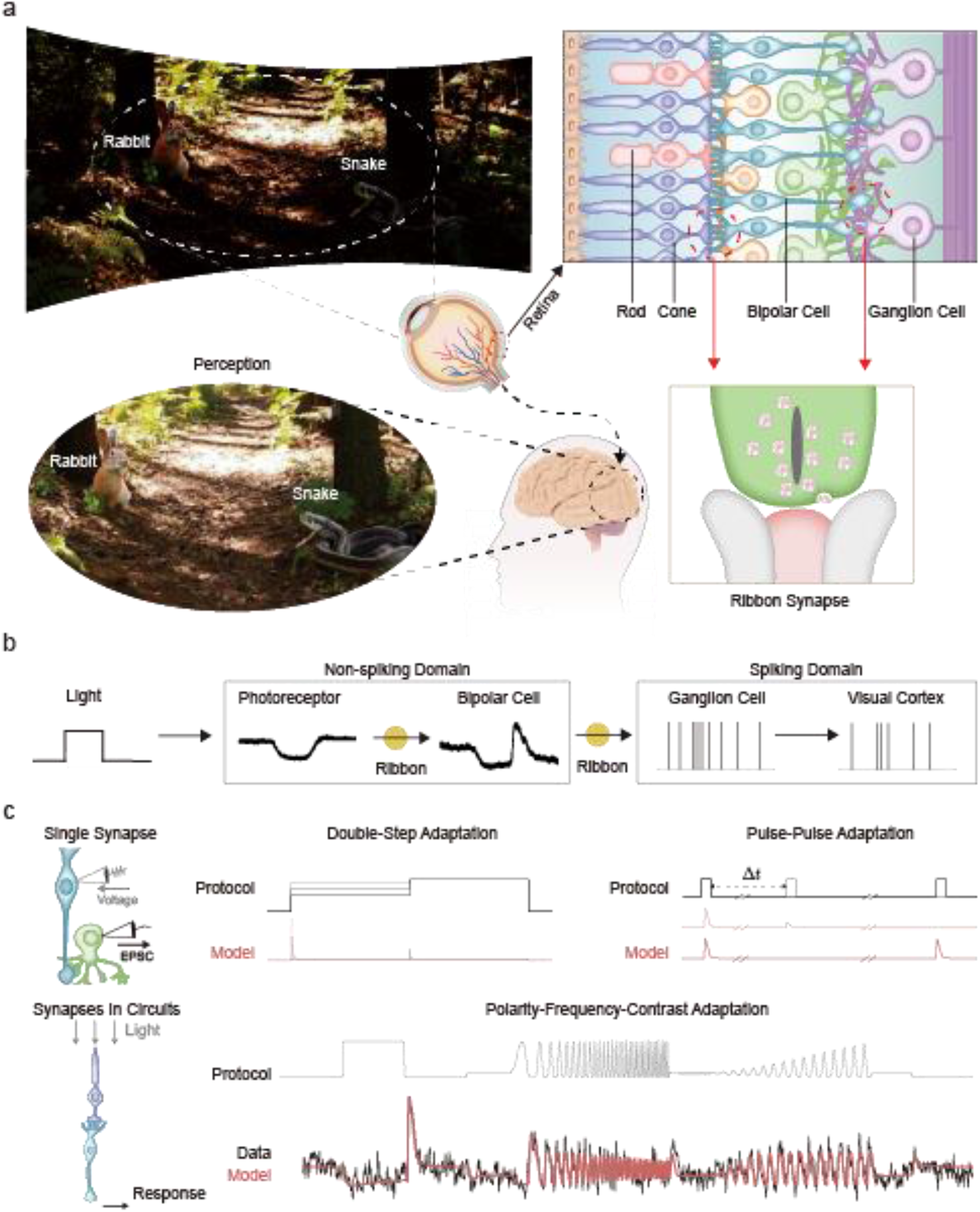
Visual adaptation in retinal ribbon synapses. (A) The retina adapts to visual contrasts through two layers of ribbon synapses. (Left) The visual system can perceive extreme contrasts in the natural environment. (Right Up) The retinal neural network is in the first stage of the visual system. Ribbon synapses are located at the terminals of photoreceptors (cone and rod) and bipolar cells. (Right Down) Diagram of the ribbon synapse with a single synaptic ribbon, an electron-dense structure illustrated as a vertical bar. Dendrites from postsynaptic neurons receive neurotransmitters released by the presynaptic terminal. (B) Due to visual adaptation, neural responses, the membrane potential in the analog domain (adapted from(12)), and the firing rate in the spiking domain are changed alongside a fixed luminance intensity. Ribbon synapses contribute to the adaptation in the analog domain. (C) Our model captures various adaptative behaviors observed at retinal ribbon synapses. (Up) At the synapse level, ribbon synapses connecting rod bipolar cells and AII amacrine cells produce transient responses at the step protocol and short-term depression at the paired-pulse protocol. (Down) At the circuit level, ribbon synapses at the terminal of bipolar cells hold intricate adaptive behaviors with the protocol involving varied frequencies and contrasts.

**Figure 2.**
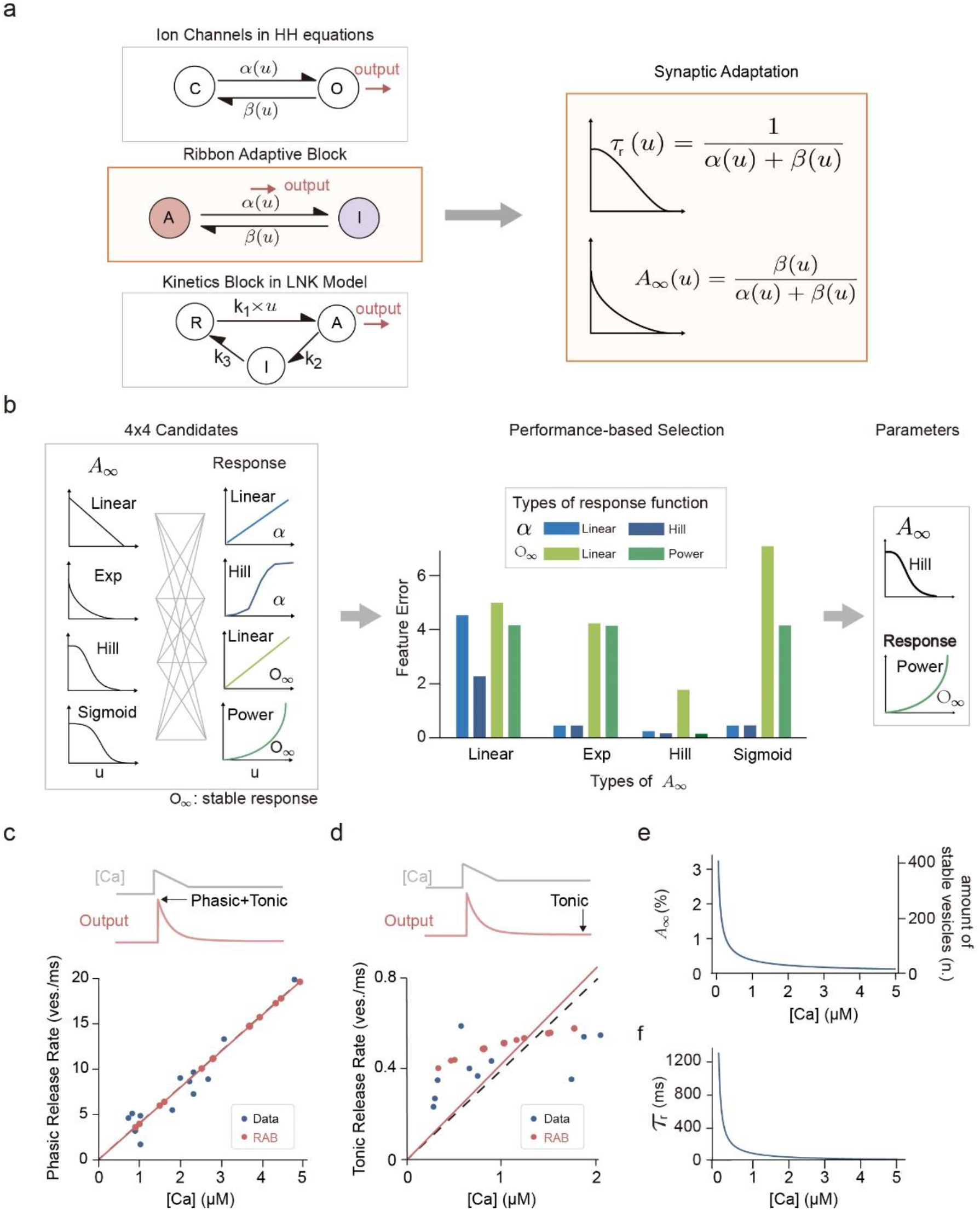
The RAB module and the pipeline of parameter selection. (A) The diagram of the RAB module and associated adaptive properties. (Left) The RAB module is a two-state transition system (middle) belonging to the family of first-order kinetics models. Compared to other kinetics models, such as the ion channel in the HH model (up) and the Kinetics block in the LNK model (down, adopted from(49)), the RAB model uses specialized transition rates and the output function. (Right) The adaptive properties, the time constant(*τ*_*r*_) and the steady-state (*A*_*∞*_) of A, are determined by transient rates between two states. (B) The pipeline of parameter section for the RAB module for salamander cone synapses. (Left) Sixteen candidates whose formulas are constrained by previous measurements of relevant biological functions. (Center) The performance of candidates compared to the experimental recordings on salamander cone synapses. (Right) The best candidate for the salamander cone is the hill function for the steady-state of the A (*A*_*∞*_) and the power function for the stable response rate (*O*_*∞*_). (C and D) The experimental protocol and the features of output for the parameter selection pipeline. (Up) A sample curve from calcium inputs and cone responses. The peak release rate is divided into a phasic component and a tonic component (C), while the stable release rate is the tonic component (D) at corresponding Ca2+ concentrations. (Down) The relationship between phasic (C) or tonic (D) release rates and Ca2+ concentrations in the experimental recordings(black) and responses simulated by the RAB module (red). (E) The steady-state of the A (*A*_*∞*_) in the optimized RAB module (the left axis) and the analogous biophysical functions (the right axis), the number of vesicles at the stable state of the synapse. (F) The adaptive time constants of the optimized RAB module for cone synapses.

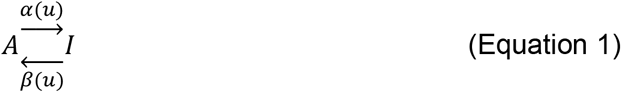

where *A* (active) and *I* (inactive) are two states conceptually related to “occupied” and “empty” vesicle sites. Accordingly, the forward transition from state *A* to state *I* is analog to the release process, and the backward transition is analog to the resupply process. As calcium signals modulate those biophysical processes, the input signal *u* is supposed to dynamically control two transitions’ rate constants (*α*(*u*), *β*(*u*)). Denoting *A* as the current value of the state *A*, the kinetics system of the RAB model can be described by the following equation:

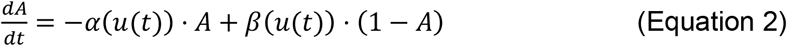

At first glance, our two-state kinetic framework shares many similarities to the classic two-state Hodgkin-Huxley (HH) formalism(34). However, the significant difference between classical state-transfer models and our model is the definition of the system’s output, where we define the transition process from **A** to **I** as the output of the RAB model, denoted as *o*(*t*). Thus, we can formulate the output as:

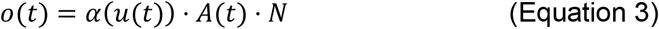

where N is a scaling factor analogous to the number of vesicle sites in the RRP.

As a two-state first-order model, the kinetics system of the RAB module can also be organized as an adaptive formulation as:

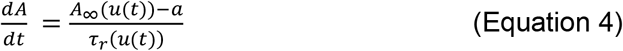

where *A*_*∞*_(*u*) denotes the steady-state of state A, analogous to available vesicles during the stable release in the RRP, and *τ*_*r*_(*u*) is the time constant of the system’s adaptive process to the steady-state. Following the similar kinetics analysis of the HH model, the adaptive properties *A*_*∞*_(*u*) and *τ*_*r*_(*u*) can be inferred from two transition rates (Fig.2A down), as

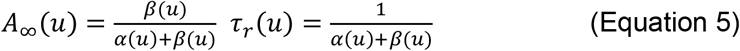

It should note that the previous linearized one-pool framework also uses a similar diagram to model the short-term depression at central synapses, which processes spike trains and derive its adaptation time constant from spikes’ firing rate(46). The RAB module is clearly different from the linearized one-pool model in terms of the definition of adaptation time constants and the models’ mathematical functions.

Till now, it can be concluded that the adaptive properties of the system are determined by the two transition processes *α*(*u*) and *β*(*u*), resembling the release and resupply process in ribbon synapses, respectively. And such biologically-constraint concepts are critical for choosing proper mathematical functions for *α*(*u*) and *β*(*u*). Below, we will elaborate on obtaining optimal parameters and mathematical functions for the RAB module.

### Pipeline for parameters selection

Equation 5 reveals the relationship between adaptive properties (*A*_*∞*_(*u*) and *τ*_*r*_(*u*)) and transition functions (*α*(*u*) and *β*(*u*)). These parameters have biophysical meanings when we map the RAB into the vesicle release/resupply process framework in the RRP (TableS1). Fortunately, with the aid of advanced techniques, we can directly measure several corresponding cellular processes, such as (1) The vesicle release rate constant(47, 48), which can be interpreted as the transition rate constant *α*(*u*), (2) the occupany rate of vesicle sites in the RRP during steady-state(16, 19), which may relate to *A*_*∞*_(*u*), and (3) the stable vesicle release rate *o*_*∞*_(*u*)(12, 32), which is determined by the Equation 6:

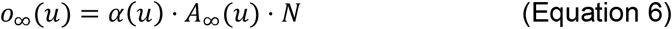

In theory, if we simultaneously measure any two functions in Equation 6, we can deduce all the rest of the functions in Equations 2 and 5. However, we usually only measure one function in a single experiment in practice. Considering these functions are diverse in different cell-type and species, we have to search for the best combinations of these functions to fit experimental curves. Therefore, we obey the following principles: first, we provide all possible combinations of biologically-constraint functions; second, we will use the brute-force approach to search for the best formulations and parameters.

Take a RAB module application for the salamander cone ribbon synapse reported in (12) as an example. We first chose 16 candidate formulations from previous experiments (Fig.2B, left). Specifically, the candidate formulations for *A*_*∞*_(*u*) include “Linear”, “Expenentional”, “Hill”, and “Sigmoid”. These common decreasing functions align with the vesicle depletion uncovered in previous experiments. Likewise, the candidate formulations for *α*(*u*) and *o*_*∞*_(*u*) are also consistent with the available data(TableS2). Next, as we only need any two functions in Equation 6 to infer the remaining functions in the model, we select either “ *A*_*∞*_(*u*) and *α*(*u*)” or “ *A*_*∞*_(*u*) and *O*_*∞*_(*u*)” combinations as the basis for inferring. Finally, at the optimization step, we test all possible combinations with optimization algorithms (Fig.2B, middle), targeting the kinetics of phasic/tonic release components in vesicle release traces (Fig.2C and 2D). Our results indicate that the best choice is the hill function for *A*_*∞*_(*u*) and the power function for *α*(*u*) (Fig2B, right). With the inferred functions, the RAB module faithfully reproduces linear relationships between phasic/tonic release rates and the input calcium signals (Fig.2C and 2D). In addition, the model predicts ribbon synapse’s available vesicles (*A*_*∞*_(*u*) ∗ *N*) at the resting is ∼418 (Fig2E), close to the estimated vesicle numbers in the original experiment (∼390)(12).

Overall, the RAB module can capture kinetics in the cone ribbon synapse’s vesicle processes by combining biologically-constrained functions and brute-force search. Interestingly, the module reveals that the adaptive time constant *τ*_*r*_(*u*) is not fixed but associated with the input strength signals, the current calcium signals ([Ca2+]) (Fig.2F), suggesting ribbon synapses might dynamically adapt to the stimulus patterns and thus exhibit complex behaviors.

### The RAB module captures adaptive behaviors in the single synapse

In this section, we aim to explore the RAB’s performance at the single-synapse level. In general, the intrinsic synaptic mechanisms drive ribbon synapses to produce enriched adaptive behaviors, such as biphasic responses(12, 16), short-term depression(17), and temporal filtering(11, 29). Yet it remains a significant challenge for a model to capture such complex behaviors fully. One general approach is allowing the model to fit these behaviors as much as possible. However, we argue that if an adaptive model can indeed grasp the intrinsic cellular mechanism, it can be trained on limited data but generalized to other “unseen” datasets. Therefore, the robustness of an adaptation model is a critical factor used to measure the extent to which the model reflects the biophysical mechanism of synapses.

We first establish a Nonlinear-RAB (N-RAB) model for a rod bipolar cell – AII amacrine cell (RBC-AII) synapse (Fig.3A)(16), composed of a static nonlinearity and an adaptive kinetics block (RAB) (Fig.3B). Specifically, the nonlinear function is taken from the calcium current-voltage curve measured in the original experiment without modifications, transforming the membrane voltage into calcium signals(16). Besides, a static linearity follows the N-RAB model, acting as a simple scalar to automatically adjust the amplitude of outputs. Hence, the remaining parameters to be tuned are all from the RAB modular, ensuring we can use the previous pipeline to obtain optimized parameters for the RBC-AII synapse.

**Figure 3.**
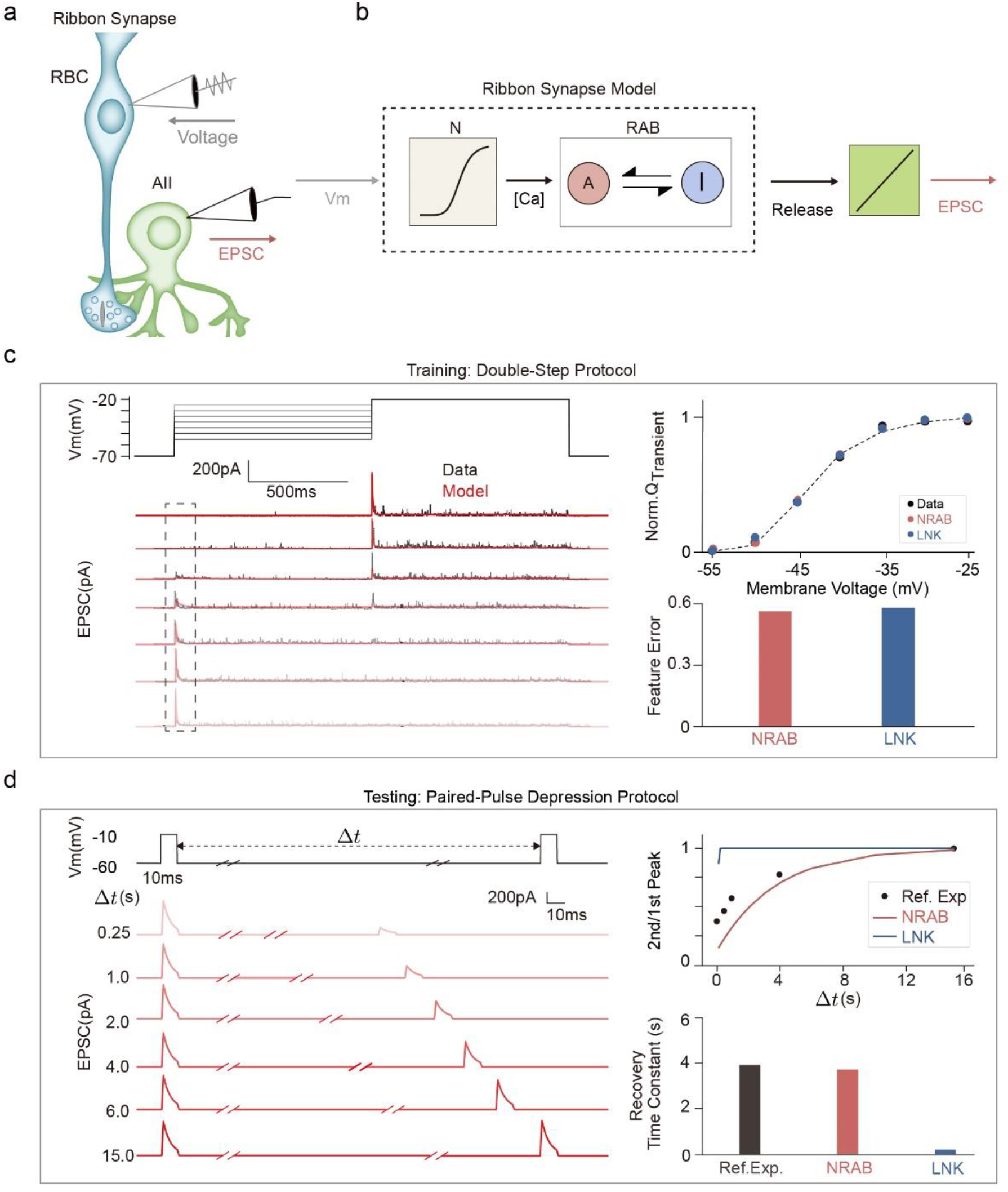
The N-RAB model captures diverse adaptive behaviors in a single synapse. (A) The experiment with the synaptic coupling of rat rod bipolar cell (RBC) and AII amacrine cell (AII) pair. It manipulates the membrane potential of RBC and records AII excitatory postsynaptic currents (EPSCs). (B) The N-RAB model of the rat RBC-AII synapse. It contains a nonlinearity to convert the membrane voltage into the Ca2+ concentration and a RAB module for synaptic adaptation. Another linear transform is appended after the RAB module to tune the amplitude of the model response. (C) The training stage of the ribbon synapse model. (Left) The double-step protocol evokes AII EPSCs. The membrane potential of the RBC synapse is stepped to a background potential for 1s, and then another 1s test step to −20mV. Response traces of experimental recordings(black, adapted from(16)) and the model simulation (red) with grayscales corresponding to input traces. (Right) A typical feature of responses for optimization and overall performances after optimization. The feature is the integrated currents over the transient component in the recordings (black) and simulated responses by the N-RAB model (red). The N-RAB model and the LNK model (blue) perform similarly on the measurement of feature errors. (D) The testing stage of the ribbon synapse model. (Left) The N-RAB model predicts AII EPSCs at the paired-pulse depression (PPD) protocol. In an individual trace, two 10ms pulses raise the membrane potential from −60mV to −10mV with a time interval Δt. Simulated responses triggered by pairs with different time intervals. (Right) The depression recovery (the peak at the 2^nd^ pulse) of responses and corresponding time constants. The recovery responses produced by the N-RAB model (red, time constant=3.9s) are similar to the reference experimental recordings (black, time constant=3.7s). In contrast, the recovery process in the LNK model (blue, time constant=0.02s) is faster than in recordings.

Next, we train the N-RAB model with postsynaptic recordings from RBC-AII synapses (Fig.3C). In the training protocol, we apply the “Double-Step” protocol to stimulate neurons: a first 1-second step-wise voltage command elevates the membrane voltage *V*_*m*_ to a range of depolarizations (−55 to −25 mV), followed by a 1-second step voltage command to −20mV (Fig.3C). The adaptive behaviors we observed are “biphasic” (transient and sustained) responses triggered by the step voltage command. Notably, the ratio of the transient currents to the maximum transient components represents how the synapse encodes temporal contrasts. Additionally, for comparison, we also trained the LNK model on the same datasets, whose adaptation block is a classical first-order state-transfer block(49). Precisely, we extract features of the responses for optimization, including transient components, amplitudes of peaks, and stable responses (see Methods). The results indicate that both N-RAB and LNK models achieve excellent performance in fitting adaptive behaviors in terms of the temporal contrast index (Fig. 3C right) and feature errors (0.56 in the LNK and 0.55 in the N-RAB.

At last, to assess the robustness of the models, we use the paired-pulse depression (PPD) protocol to test the short-term plasticity. Unlike the previous Double-Step protocol, the PPD protocol uses two pulses to trigger transient responses(Fig.3D). The 2^nd^ response is depressed if the interval between two pulses is short. Thus, the relationship between intervals and synapses’ 2nd responses is essential for measuring recovery time constant, an index for short-term depression. We first apply the protocol to the trained N-RAB model and find the model produces an intense depression in the 2^nd^ response. We then use an exponential function to measure the recovery time constant (τ=3.7s), which is consistent with another measurement (τ=3.9s) with the same protocol in RBC-AII synapses (Fig.3D right)(50). By contrast, the LNK model’s recovery time constant is fairly fast (τ=0.02s), unable to reproduce the depression responses during the PPD protocol.

In summary, the experiment results suggest that the N-RAB model is highly robust for synaptic adaptation. It captures the intrinsic mechanism through data-fitting on the training data and generates reasonable responses even to unseen protocols. We expect that the adaptive properties revealed by the N-RAB model may further deepen our understanding of the biophysical mechanisms of adaptation at the synapse level.

### How the RAB module produces adaptive behaviors

Why the RAB module shows good generalization to various stimulus patterns? We first examined the curves for the adaptive parameters *A*_*∞*_(*u*) and *τ*_*r*_(*u*). Interestingly, both *A*_*∞*_(*u*) and *τ*_*r*_(*u*) inferred from the “Double-Step” protocol are similar to inverse S-shape curves, which change rapidly between −60 mV and −40 mV. From the view of HH formalism, such kinetic curves look like an inactivation gate of ion channels. Surprisingly, the time constant *τ*_*r*_(*u*) of the RAB module can switch from seconds to milliseconds during the ∼20 mv transition. By comparison, the inactivation gate of typical ion channels typically changes from tens of milliseconds to milliseconds during the same transition(34). Therefore, the dynamic range of the RAB time constant is ∼10-100 folds larger than those of typical inactivation gates, which enables RAB to fastly adapt to stimuli at more depolarized *V*_*m*_ (>-40 mV) and have strong capacitance to remember the previous state at more hyperpolarized *V*_*m*_ (<-60 mV).

Inner dynamics suggest that different membrane potentials drive fast or slow adaptation in the training and testing protocols. During the first Double-Step protocol, the membrane stays at depolarized voltages where the time constant *τ*_*r*_(*u*) is less than 50ms(Fig.4A left) so that the state variable A rapidly decreases along with *A*_*∞*_(*u*), approaching an empty state(Fig.4B left). During the second PPD protocol, the first pulse elevates the membrane voltage *V*_*m*_ to − 10 mv, leading to a dramatic decrease in *A*_*∞*_(*u*), similar to the transient response in the Double-Step protocol. However, the recovery of a depressed synapse occurs at a hyperpolarized *V*_*m*_, −60mV, which yields a large time constant *τ*_*r*_(*u*) (∼3-4 seconds, Fig.4A right) and determines the slow recovery process from depression(Fig.4B right). Thus, if a second pulse arrives early, the slow time constant *τ*_*r*_(*u*) will prevent A from regaining the previous state, resulting in a “depression” in response to the second pulse (FigS.3). Hence, the inverse S-shape curves of adaptive time constants *τ*_*r*_(*u*) empower a RAB module producing behaviors with distinct adaptive speeds, matching experimental recording at different protocols.

**Figure 4.**
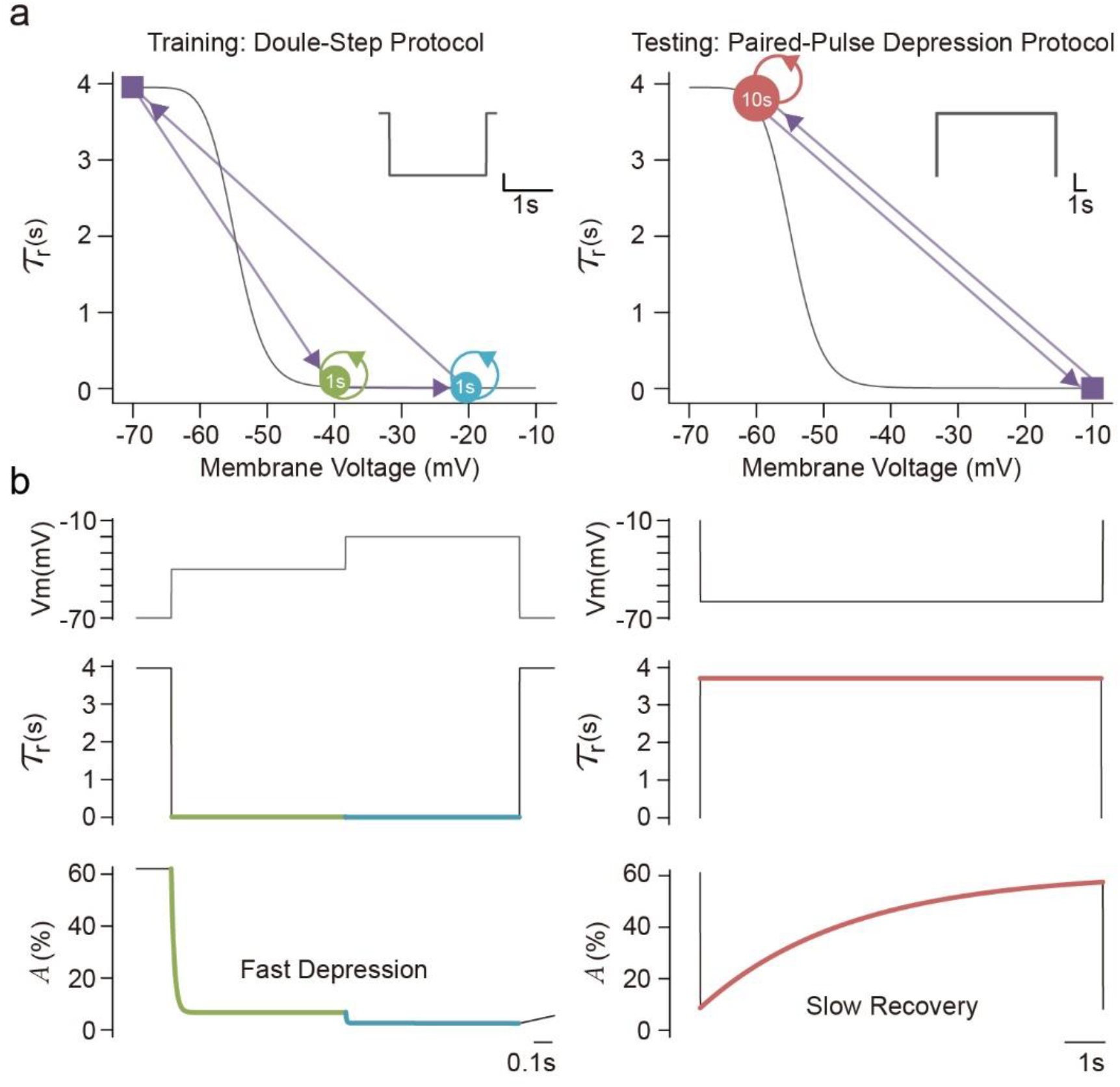
Inner dynamics of the RAB module. (A) Adaptive time constants at two protocols. (left) The double-step protocol triggers transient responses at high potentials with small time constants (<50ms). (right) The PPD protocol recovers the active state A at a low potential with a large time constant (3.7s). (B) Inner variables during the fast depression at the double-step protocol (left) and the slow recovery at the PPD protocol (right). From up to down: the input profile of the membrane potential, adaptive time constants, and values of the active state A alongside the stimuli.

To conclude, the dynamic adaptive parameters *A*_*∞*_(*u*) and *τ*_*r*_(*u*) inferred from the training protocol endow the N-RAB model with remarkable generalization ability, suggesting the RAB modular is not simply data-fitting but instead captures intrinsic biophysical mechanisms of ribbon synapses.

### The RAB module captures adaptive behaviors in circuits

Diverse biophysical mechanisms in circuits accomplish visual adaptation in the retina, typically in phototransduction, local feedback, lateral inhibition, ion channels, and synaptic transmission(6). Furthermore, neural activities of bipolar cells(BC) and ganglion cells (GC) subtypes are distinct from other subtypes, suggesting synapse-related processes behind cells are type-specific(35, 51-54). Therefore, components in the retinal circuit obscure the critical role of ribbon synapses in visual adaptation. To reveal underlying mechanisms in ribbon-related visual adaptation, we pick the two-photo imaging data recorded with glutamatergic outputs of mouse BCs(35), which eliminates the effects of interneurons and the spatial adaptation by pharmacological manipulations(+glycine receptor block) and the local stimulus protocol(100μm diameter)(Fig.5A). Notably, BCs are categorized into multiple subtypes, including three Off-types(*C*_1_, *C*_2_, *C*_3*a*_) and three On-types(*C*5*o, C*_*X*_, *C*_7_). Distinct morphological and electrophysical properties characterize these subtypes, yet their functional diversity exhibited during visual adaptation is not fully explored(55). Fortunately, the chosen data was obtained from large-scale detectings of synaptic activities at the level of individual terminals, providing reliable recordings covering all these subtypes of BCs. Moreover, the input protocol used a widely used ‘chirp’ stimuli(56, 57), comprising varied frequencies and contrasts and triggering worthy adaptive behaviors for capturing by theoretical models(Fig.5C). Previous work has established an elegant theoretical method to infer the activities of ion channels in BC’s dendrites(58). Based on multi-compartment models, this approach inevitably has high computational complexity. Plus, it didn’t check synaptic adaptations in visual processing. Thus, we develop a parsimonious model with RAB blocks to address synapses in retinal circuits.

**Figure 5.**
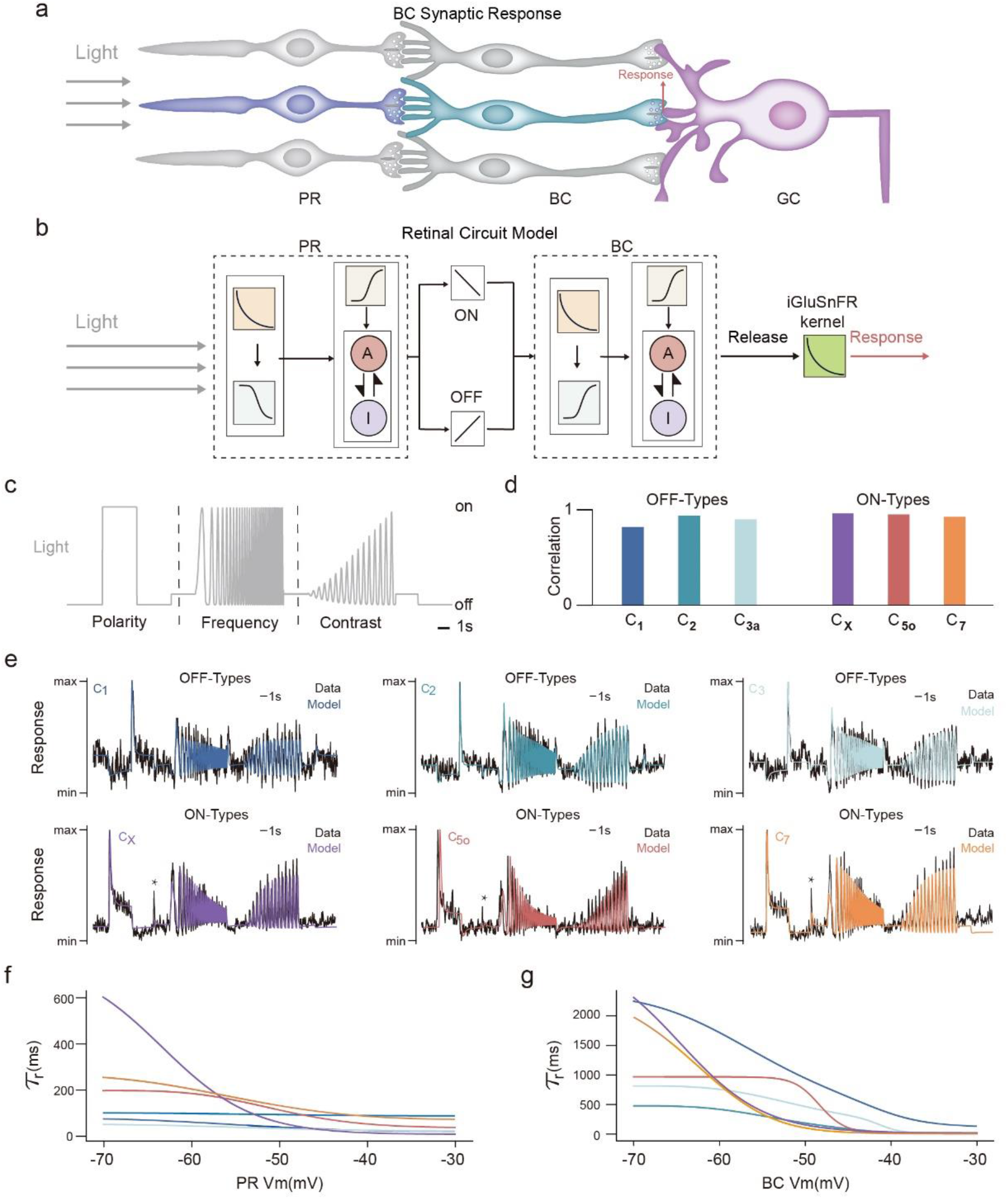
The hLNS model captures synaptic behaviors in the Photoreceptor (PR) – Bipolar Cell (BC) – Ganglion Cell (GC) circuit. (A) The experiment on mouse bipolar cells. It changes the light intensity and records vesicle release rates at BC synapses. (B) Schematic of the hLNS model of the PR-BC-GC circuit. From left to right: a photoreceptor cell with synapses, a linear transform to control the polarity of synaptic responses, and a bipolar cell with synapses. The model uses the LN block to transmit the inputs to the membrane voltage and the N-RAB block to mimic the synaptic activity. (C) The light stimuli in the ‘chirp’ protocol. It uses a step to test the polarity of BCs, followed by a stage with temporal frequencies from 1Hz to 10Hz and another stage with contrasts from 0% to 100%. (D) Correlations of the experimental recordings and the responses simulated by the hLNS model in circuits with different BC subtypes. (E) Light-evoked traces for six BC subtypes (three Off-types and three On-types). Overlaid are responses from experimental recordings (black) and model-simulated (colored). (F and G) Adaptive time constants of PR-BC synapses (F) and BC-GC synapses (G) in six subtype circuits.

Here, we establish a model with two layers of cells to describe the photoreceptor(PR)-bipolar cell(BC) circuit. We mainly follow classical models for BC’s synaptic activities, the Linear-Nonlinear(LN)-Synapse architecture, which contains an LN block to convert the light signal to the membrane potential or calcium signals, and a synaptic block to produce responses(27, 29). However, these models ignored unobserved hidden adaptations in PR-BC synapses. A standard approach to reconstructing unobserved behaviors in multi-layer circuits is cascaded linear-nonlinear (LN-LN) models(25, 26), using LN units to represent individual neurons so that connections among units can directly map onto circuit anatomy. Here, we propose the hierarchy Linear-Nonlinear-Synapse (hLNS) model, combining the LN-LN architecture with synaptic adaptive blocks to mimic the two layers of synaptic adaptations in the PR-BC circuit (Fig.5B). In particular, each cell in an hLNS model contains one LN block to generate the membrane potential from the input signals (light or upstream signals) and one N-RAB block to generate postsynaptic currents or glutamate releases of the cell. Two cells in the circuit are connected with a linear function to model the polarity of signals after postsynaptic receptors. We use the best-fitted formulations of cone synapse in (Fig.2A) for both PR and BC synapses. Additionally, we add necessary biophysical constraints of cell membrane potentials during models’ optimization to ensure the rationality of the neural activities (see Methods).

The optimized hLNS model reproduces all types of BCs’ responses in separated stages (Fig.5E): 1) At the polarity stage, the model response reaches the maximum peak value and shows strong polarity. 2) At the frequency-varied(0.5 to 8Hz) stage, peaks decreased with the incremental frequency, mainly due to the deprivation of the active state in the 2^nd^ RAB block(FigS.5A). 3) At the contrast-varied stage, the elevated contrast triggers high membrane potentials in the 2^nd^ layer so that the model produces gradually increased outputs(FigS.5A). Overall, the hLNS model successfully captures the complex adaptive behavior at the ‘chirp’ light stimuli (correlation coefficient between the model and the response is 89 ± 5%, mean ± SEM, Fig. 5D).

Furthermore, the hLNS model infers the diversity of synaptic adaptation in neural circuits. First, we find that in the circuit, the 1^st^ layer synapse(Fig.5F) has a larger adaptive time constant at the resting potential than the 2nd layer(Fig.5G), indicating the depression recovery of BC synapses is slower than PR-BC synapses. Experimental results among species are consistent with our findings, as BC synapses’ recovery time constant is several seconds(2s in goldfish(59), 3.9s in rat(50)). In contrast, PR-BC synapses’ are hundreds of milliseconds with postsynaptic recordings (100ms∼800ms in squirrel(11, 60), 300∼500ms in the salamander(12, 17)). Second, the PR-BC connection in the *C*_*X*_ circuit has a significantly large adaptive time constant at the resting potential (red line in Fig.5F). In neuroanatomy, the *C*_*X*_ is morphologically identified as a “giant” BC with a sparsely branched dendrite(36), with non-invaginating contacts to cones while other subtypes of On-type Cone BCs are invaginating contacts(Fig.6A)(37). Nevertheless, its electrophysiological properties and specific role in visual processing remain unclear. In the next section, we will investigate the type-specific adaptation at the PR-*C*_*X*_ synaptic connection.

**Figure 6.**
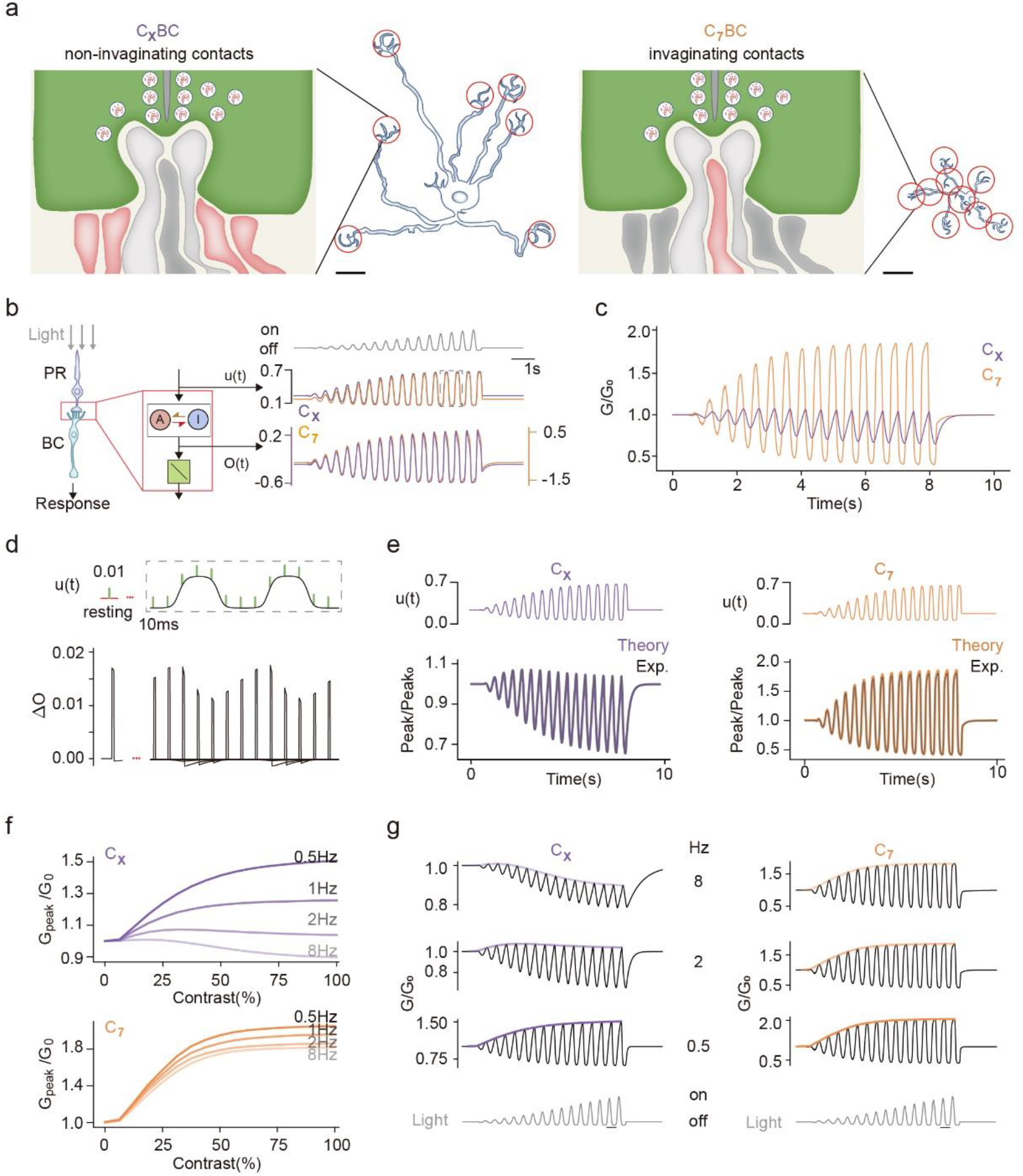
Frequency-sensitive gain-control in the PR-*C*_*X*_ BC synapse. (A) Morphologies of two BC subtypes and their contacts to photoreceptors. Compared with the *C*_7_ BC, the *C*_*X*_ BC has a larger dendrite field and non-invaginating contacts(adapted from(36, 55)). (B) The input (u(t)) and the output (O(t)) of the PR-BC synapse in two circuits (*C*_*X*_ and *C*_7_) are similar. (C) The theoretical gain (Equation 7) of the PR-BC synapse in the *C*_*X*_ circuit (yellow) and *C*_7_ circuit (purple) at the contrast-varied stage. Peak values of the gain in the *C*_*X*_ circuit decreases with the increase of the contrast. Values are normalized by the gain before the stage. (D) The experimental method to measure the gain of the RAB module in the PR-BC synapse on the *C*_*X*_ circuit. It delivers 10ms flashes to the input stage (u(t)) and calculates changed responses (ΔO) on the output. (E) Theoretical analysis results of the gain match the experimentally measured changed responses of the PR-BC synapse in the *C*_*X*_ circuit (left) and the *C*_7_ circuit (right). (F) The PR-BC synapse in *C*_*X*_ circuit (up) shows frequency-sensitive gain-control patterns compared to the *C*_7_ circuit (down) whose are unchanged with varied frequencies. (G) Gain traces in the varied-frequency protocol. This protocol changes the temporal frequency of the contrast step from 0.5Hz to 2Hz (bottom, gray). The gain of the PR-BC synapse in the *C*_*X*_ circuit is suppressed at high frequency (left), while the gain in the *C*_7_ circuit is unchanging (right). Curves illustrate the trends of peak gain with evaluated contrasts.

### High-frequency triggered depression of gain in the *C*_*X*_ PR-BC synapse

The complex input protocol prohibits the understanding of how type-specific synaptic adaptation in the *C*_*X*_ PR-BC synapse contributes to visual adaptations. Taking the *C*_7_ circuit for comparison, at the contrast-varied protocol (the 3rd stage of the ‘chirp’ stimuli), inputs and outputs of the 1st layer RAB modules in two circuits are too similar to distinguish their adaptations(Fig.6B). Such similarity between *C*_7_ and *C*_*X*_ adaptive responses are supported by an electrophysical experiment(54). To distinguish intrinsic adaptative mechanisms in two RAB modules, we introduce the gain, the amplitude of the response to an input signal, as a measure of visual adaptive properties. Generally, the visual adaptation is interpreted as a gain control strategy, with dynamic gain depending on the historical stimuli(2). Previous work measured the gain control in a retinal cell(10) or an adaptive model(49) through experiments. Here we present an analytical approach to obtaining the gain of the RAB module. Following the classical definition(49), we define gain as the change in response (*Δo*) caused by a small perturbation in the input (*Δ*_*u*_). Thus the gain at time t (*G*(*t*)) can be written as

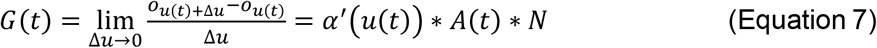

where *α*(*u*) is the transition rate constant for the given input *u, A*(*t*) is the current active state, and *N* is the scaling factor of the RAB module(see Method). Notably, the formulation of the gain *G*(*t*) (Eq.7), proportional to the derivative of the transition rate constant *α*(*u*) instead of the value of *α*(*u*), differentiating it from the previous formulation in the LNK model(49) due to the specialized output function in the RAB module.

By analyzing the 1^st^ layer of RAB blocks in the *C*_*X*_ and *C*_7_ circuits, we observe two distinct gain-control strategies at the 3^rd^ stage of the ‘chirp’ protocol, where contrasts are elevated during stimulation. The gain in the *C*_*X*_’s block is gradually decreased with the rise of the contrast, while the *C*_7_’s block maintains the gain at a higher level even at 100% contrast (Fig.6C). To verify these analytical results, we use an experimental approach to observe the synaptic gain by delivering brief flashes of *u* at different timing points (Fig.6D). After subtracted by the original response curve and normalized by the amplitude at the resting state, the peak values of flash-triggered responses precisely fall on the instantaneous gain curve, verifying the theoretical finding of the depressed gain in the RAB module representing adaptive functions of *C*_*X*_ PR-BC synapse(Fig.6E).

To unveil the underlying mechanism of the gain depression, we examine inner dynamics in RAB blocks and find that the deprivation of the active state is the leading cause (FigS.6B). During the stimulation with varied contrasts, the temporal frequency of the input is fixed at 2Hz. In this situation, the interval between two peaks(500ms) is an intermediate value between adaptive time constants at low potentials in two synapses. Consequently, the larger time constant(∼600ms) in the *C*_*X*_ RAB block prevents the active state A from recovery, while the relatively shorter time constant(∼300ms) in the *C*_7_’s block helps its active state almost return to previous peaks (FigS.6C&D). Therefore, the 2Hz stimuli trigger distinct gain-control patterns in two RAB blocks. Next, to verify this frequency-based analysis, we shift the light’s frequency from 0.5Hz to 8Hz and find the gain decreased significantly in the *C*_*X*_ block(Fig.6F&G). By comparison, the *C*_7_ block’s gain pattern is unchanged with varied frequencies. Besides, such frequency-sensitive phenomena do not appear in the synaptic responses (FigS.6F&G). These results indicate that the PR-*C*_*X*_BC synapse is more resistant to perturbations in fast-changed and high-contrast inputs, which might appear in the natural environment.

In conclusion, we theoretically analyze the gain of RAB blocks with the frequency- and contrast-varied protocol and find the special gain control strategy of PR-BC synapse in Cx circuits using high-frequency and high-contrast stimulus. In contrast to previous experimental protocols, which varied frequency or contrast in separated stages, we provide an approach to access the synaptic adaptation with more complex protocols. We hope our tools, the hLNS model for capturing adaptation and the gain for quantifying adaptation, can help researchers understand the complicated role of ribbon synapses in visual adaptation.

## Discussion

This study provides the RAB module for synaptic adaptation in ribbon synapses and the hLNS framework for retinal circuits. We established a series of models that successfully captured transient vesicle releases in salamander cone synapses(12), postsynaptic responses in rat rod bipolar cell synapses(11), and ‘polarity-frequency-contrast’ responses in mouse cone bipolar cells with light(35). At the synapse level, the RAB module and the extended N-RAB model show a high degree of generalizability in inferring intrinsic adaptive properties. Besides, we found that the formulation of transient rates in the RAB block determines the N-RAB model’s generalization. Hence, we credited the good performance of our model to its biophysical constrained transient rates compared to constants or handcrafted ones in previous models. At the circuit level, we illustrated the diverse adaptive properties of two-layer ribbon synapses in the PR-BC-GC circuit and found the PR-*C*_*X*_ BC synapse has a larger adaptive time constant than other PR-BC synapses in On-type circuits. To reveal the role of adaptive time constant, we investigated the gain-control in the PR-*C*_*X*_ BC synapse and found that its relationship between gain and contrast is sensitive to the frequency of stimulation. In conclusion, the RAB model bridges the gap between biological mechanisms and synaptic adaptations, allowing us to uncover the intrinsic adaptive properties inside retinal circuits.

### Biological constraints in models

At the synapse level, we modeled processes of vesicle release and resupply in the RRP and then added constraints on the parameters with the electrophysical properties of ribbon synapses. These parameters vary across different cell species, yet it remains a challenge, given the limitation of current experimental techniques, to launch an extensive study over diverse types of synapses and fill all the blanks in parameter measurements. Therefore, we provide a pipeline to theoretically determine all these parameters and formulations based on current experimental results (Fig.2B). We show that the cell’s chosen formulations are consistent with the cell type in the experiment. For instance, the power stable response rate in the model for the salamander cone synapse(Fig.2) comes from an experiment on the same species and the cell type(32). Similarly, the hill release constants for the rat rod bipolar cell model (FigS.1A) are from an experiment on goldfish bipolar cells(48).

At the circuit level, we introduce membrane potential constraints to the hLNS model. The retinal circuit experiment only measured synaptic responses in the second synaptic layer but lacked measurements of internal neural activities(35). Hence, we apply several constraints to ensure the rationality of the internal variables. First, the potential of each cell must be negative (<0mV), as neurons in this circuit do not generate actional potentials(8). Second, we ensure that the membrane potential of dark-adapted cells is close to experimentally measured values, such as −46mV for cones(61), −43mV for off-type BCs(62), and −66mV for on-type BCs(63). Besides, nonlinearities in all N-RAB blocks are directly adopted from the calcium voltage-current curve in experimental measurements. These constraints successfully narrow down the space for parametric searching in the case of the private line circuit with local light stimuli.

### Related models

Retinal synapse models typically focus on the process of vesicle movements at presynaptic terminals. Mechanistic models use multiple pools to detailedly describe the life cycle of vesicles from endocytosis to exocytosis(27, 29, 30). Another popular approach is to simplify the complicated life cycle into vesicle release and resupply of a single pool, the RRP(12, 16, 58, 64-66). However, mechanisms inside ribbon synapses, such as the vesicle attachment to the synaptic ribbon(31), and the diffusion of vesicles along the surface of the synaptic ribbon(33), are not well-studied. Therefore, ribbon synapse models face challenges in determining the parameters of vesicle movement rates. To address this, these models use constants or handcrafted formulations, which are prior designs in the models and may not be flexible enough to capture some critical features in the synaptic adaptation(TableS3). In contrast, the RAB model is analogous to the single-pool framework but uses formulations constrained by experimental measurements, reducing the model uncertainty and improving model accuracy for predicting adaptive behaviors.

At the circuit level, ribbon synapses are often presented as nonlinear functions in the Linear-Nonlinear(LN)-based models, such as half-wave rectifiers(67) or cumulative Gaussian functions(68). These models elegantly generate adaptive behaviors through circuit-based mechanisms, such as local feedback(25, 51), lateral inhibitions(69), or gap junctions(70), but to some extent ignore the intrinsic adaptative properties of ribbon synapses. Recent studies have attempted to identify the role of synapses in circuit-related adaptations by adding mechanistic ribbon synapse models(28) or phenomenal kinetics blocks into the LN-based model(24, 49). In those models with complex kinetics systems, linking ribbon-related functions to adaptations is measured experimentally rather than theoretically. Furthermore, we find these models fit specific data but generalize poorly to unseen stimuli, especially the recovery from depression at paired pulses(FigS3&FigS4). In contrast, our models highlight the greater power of generalization through the specific design of two-state kinetics blocks, enabling us to understand synaptic adaptation and capture properties with limited experimental recordings.

### Comparisons with experimental findings

Simulations of models with RAB modules reveal that adaptive time constants of ribbon synapses are cell-type-specific and well supported by direct measurements as well as anatomical and electrophysiological evidence. For instance, the adaptive time constants for most of the RAB modules at the depolarized membrane potential(−10mV) are located in a small range (1ms∼10ms), supported by the fast decay in transient responses on various types and species((50, 71, 72), systematically reviewed by(7, 9, 73)). In contrast, at a hyperpolarized membrane potential(−70mV), adaptive time constants range from ∼100ms (the mouse Photoreceptor(PR) - OFF-type Cone Bipolar Cell(CBC) synapse) to ∼4s (the rat Rod Bipolar Cell – AII Amacrine Cell synapse), supported by measured short-term depressions among retinal synapses(17, 59, 74). At the circuit level, we predict that adaptive time constants of PR - ON-type BC synapses vary from 200ms in the PR-C5o synapse to 600ms in the PR-*C*_*X*_ synapse, while those synapses share upstreaming neurons(55). Experiments haven’t measured these synapses’ short-term depression in the mouse retina. Indirect supporting evidence is from PPD experiments on ground squirrel cone - Off-type CBC synapses’ presynaptic terminals, which find that adaptive time constants at −70mV also vary from 100ms to 800ms, depending on the BC types(60).

Additionally, by fitting synaptic responses of BCs, the hLNS model predicts that the PR-*C*_*X*_ BC synapse has a larger adaptive time constant at hyperpolarized membrane potential than other ON-type PR-CBC synapses suggesting that the PR-*C*_*X*_ BC synapse has a relatively slow recovery at the paired-pulse depression. Anatomical and electrophysiological findings support this prediction: First, the *C*_*X*_ BC mainly makes atypical non-invaginating contacts to M-cones instead of invaginating contacts in other ON-type CBCs(37). Then, non-invaginating contacts are larger recovery time constants at PPD than invaginating contacts, even if they share the same presynaptic pedicle(11). In conclusion, the relationship between different contact types and synaptic recovery time constants revealed by the hLNS model is consistent with the experimental findings in PR-*C*_*X*_ BC synapses.

### Gain control in the RAB block

The sensory system needs to balance its adaptation to historical stimuli with its sensitivity to future stimuli to encode a wide temporal range of input signals. ‘Gain’, the amplitude of response to input signals, is used to identify neural adaptive behaviors and describe how the system controls its sensitivity to small perturbations(2). The gain of retinal neurons’ responses decreases at high contrast, suggesting the neurons become insensitive to stimuli. On the contrary, the gain increases at low contrast so that neurons can reliably detect small environmental changes(38). Such “gain control” at different levels of input contrast was also found in ribbon synapses of mouse rod bipolar cells by introducing noise with different standard deviations into the membrane potential(18).

Compared to the previous study(18), we alternated the gain measurement protocol to ensure our results’ biological interpretability. First, we measured the gain by adding perturbations on the input of the RAB module, simulating internal noises in the calcium channels(75) instead of membrane potential fluctuations. Second, the membrane potential in our experiments oscillates between −70mV and −45mV (Fig.6A). In contrast, the membrane potential in(18) is set around −51mV or −45mV with a small standard deviation (1.5mV or 4.5mV). The wide-range oscillation pattern can clearly describe cell membrane potentials’ dynamics in the natural environment. Interestingly, we discovered new properties of the PR-*C*_*X*_ BC synapses’ gain control strategy with our biologically realistic protocol. We found that given oscillating inputs, the PR-*C*_*X*_ BC synapse adjusts its gain according to the frequency of stimuli (Fig.6F). Our predictions suggest that the gain-control in ribbon synapses is more complex than the ‘reduced gain at high contrast’ rule in previous experiments.

### The Extensibility and Future development

The flexibility in our theoretical approach provides the high extensibility of the RAB module so that we can address various biophysical mechanisms introduced by experimental protocols. For instance, experiments used different stimuli to trigger synaptic activity, such as calcium signals(Fig.2), membrane potential(Fig.3), and light(Fig.5). Theoretical models have been customized depending on experiments, combining RAB blocks as the core of the synaptic adaptation and other components to obtain additional related mechanisms. Specifically, at the subcellular level, we add a nonlinearity to represent the activation of L-type Voltage-Gated Calcium Channels so the model can receive the membrane voltage as the input(similar to(29)). At the cellular level, we add an LN block before the synapse to represent how the neuron converts the inputs into the membrane potential (similar to(22)). Similarly, the RAB module can be plugged into various experiments without modifying its kinetics system. Future development should add more components to represent relevant mechanisms in synaptic adaptations. At the subcellular level, the N-RAB model’s nonlinearity ignores several mechanisms that modulate the calcium signal for vesicle releases, such as the Calcium-Induced Calcium Release (CICR(76)), the expansion of calcium microdomains(77), and the variation of V-I curves in individual active zones(78). Thus, we can replace this nonlinearity with a more biophysical model for calcium signals, such as the HH-like equation for the calcium channel, the calcium buffer model with depths (64), and the kinetics model for the CICR(79), or a combination. At the cellular level, the role of horizontal cells, which can provide inhibitory feedback to photoreceptor terminals(80), is ignored in our hLNS model. For more general circuit models, the circuit model must contain neurons and neural connections, representing more complicated circuits(4, 6, 81). Furthermore, an ideal circuit model needs to receive spatial inputs instead of a single value in our model to explain experiments with spatiotemporal stimuli, such as visual pathway interactions (rod-cone(82), center-surround(68), etc.).

## Methods

### Code availability

The source code of models and experiments in this work will be available on GitHub and ModelDB.

### Details of the RAB module

The RAB module is a two-state first kinetics order model. States and rate constants are defined as

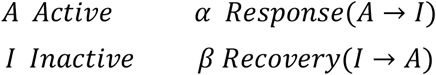

The inputs, *u*(*t*), decides transient rate constants (*α*(*u*(*t*)) *and β*(*u*(*t*))) along the time. The inactive state is derived from

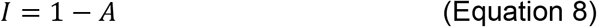

Then the active state is simulated by Equation 2. The model’s output is the transition rate *A* → *I* (Equation 3).

To get the steady-state of A and the time constant of the system, we first let the right of Equation 4 equal 0 and derive the steady-state (Equation 5). Then Equation 4 was transformed into

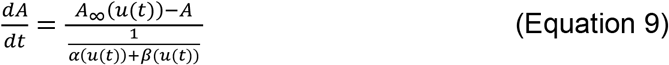

Thus, we got the time constant of the system (Equation 5). The active state A is a temporal filter for history stimuli *u*(*t*), and its time constant depends on the strength of input, as

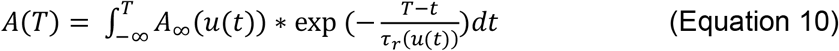

### Formulations of parameters in the RAB module

The two-state kinetics system in the RAB module is analogous to the vesicle release and resupply processes in the RRP. In a ribbon synapse, the input for these processes is the Ca2+ concentration, and the output is the vesicle release rate. Thus, several variables in the RAB model can be mapped into experimental measurements in biophysical processes (TableS1). In theory, once *A*_*∞*_ and *α* are given, we can derive the value of *β* and *τ*_*r*_, as

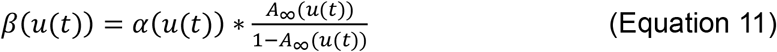

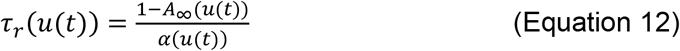

In our applications, we determined the formulation of variables by existing measurements on the stable occupancy rate of vesicles (analog to *A*_*∞*_) and the vesicle release rate constant (analog to *α*). Besides direct measurements of release rates, stable release rates (analog to *O*_*∞*_) can also provide a reference for the formulation of *α*, as

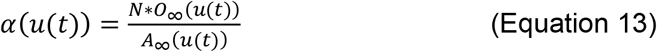

The detailed formulations of candidates are summarized in Table S2. In addition, early single-pool models for ribbon synapses can be viewed as the RAB model with different formulations of rate constants *α* and *β* (Table S3).

### The Linear-Nonlinear block and the calcium nonlinearity

We used an LN block to convert the stimuli for a cell (*x*(*t*)) into the membrane potential *V*_*m*_(*t*) and then a nonlinearity to convert the membrane potential *V*_*m*_(*t*) into the Ca2+ concentration *u*(*t*).

In an LN block, the input *x*(*t*) is passed through a linear temporal filter *L*(*x*(*t*)), a static nonlinearity *N*_*LNS*_(L), and a fixed linear transform *V*(*N*_*LNS*_), as

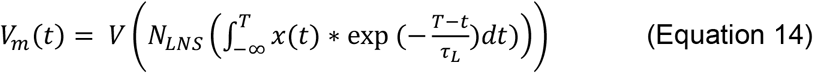

where *τL* controls the temporal sensitivity of the linear filter. After the linear filter *L*(*x*(*t*)), the nonlinear function *N*_*LNS*_ transforms the shape of response, as

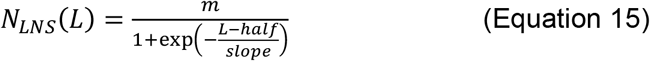

where *m, half*, and *slope* are parameters for optimization. At last, *V* converts the value of response into the membrane potential, as

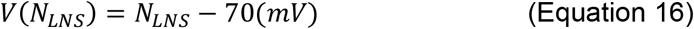

In an N-RAB model or block, the nonlinearity *N*(*V*_*m*_(*t*)) transforms the membrane voltage *V*_*m*_(*t*) into the ca2+ concentration *u*(*t*). In cases where the calcium V-I curve is not measured in the original experiment, we adapt the I-V curve of Cav1.4 in salamander cones(78), as

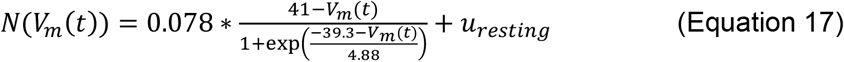

where *u*_*resting*_ = 0.05 presents the resting Ca2+ concentration in terminals(12). In the model for rat rod bipolar cell (Fig.3), we adopt the measured V-I curve (*I*_*Ca*_ in Fig.2b of(16), as

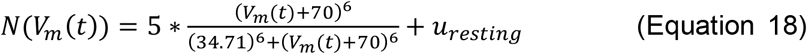

### Constraints for parameters and variables

For the RAB module, we add several constraints for *A*_*∞*_(*u*), *τ*_*r*_(*u*) and *O*_*∞*_(*u*). To mimic the vesicle depletion in ribbon synapses, *A*_*∞*_(*u*) should be: a) 0 *≤ A*_*∞*_(*u*) *≤* 1. b) nonincreasing function. c) *A*_*∞*_(0) = 1, expect the sigmoid case, as the value of the sigmoid function < 1. Meanwhile, 1*ms* < *τ*_*r*_(*u*) < 10*s*, to ensure the module finds the relatively fast adaptation in synapses instead of other slow adaptations in circuits(6). For the stable response *O*_*∞*_(*u*) must increase with the rise of the inputs. Besides, to avoid delicate and vulnerable parameters, the amplitude of raw output should be larger than 1e-3.

In addition to parameters, we add several constraints for the input of the RAB module *u* and the membrane potential *V*_*m*_. The value range for *u* is (*u*_*resting*_, 5*μm*]. The value of membrane voltage *V*_*m*_(*t*) is constrained in the range [-70mV, 0mV). In the hLNS model, We restrict cells’ membrane potential at dark (*V*_*dark*_). Specifically,

1. The 1^st^ layer of LN block’s *V*_*dark*_ should be around −46mV (from the macaque retina(61)).
2. The 2^nd^ layer of LN block’s *V*_*dark*_ in the off-type circuit should be around − 43mV(from the mouse retina(62)).
3. The 2^nd^ layer of LN block’s *V*_*dark*_ in the on-type circuit should be around − 66mV(from the squirrel retina(63)).

The tolerate range is ±5mV for all blocks.

### The experiment on the salamander cone synapse

All experimental protocols and referenced recordings come from(12). In the original paper, calcium concentration traces ([Ca], *u* in the model) are measured with a large timestep(200ms), which means we cannot directly use them in the simulation. We artificially generated 20 calcium traces following the description in the experiment. For each trace,

1. *u* is resting at *u*_*resting*_ = 0.05*μM*.
2. At t=200ms, *u* is lifted to *U*_*top*_.
3. During 200ms∼700ms, *u* is linearly decreasing to *U*_*down*_.
4. After 700ms, *u* is kept at *U*_*down*_ until the trace ends at 1200ms. In a single trace, *U*_*top*_ ∈ [0.7*μM*, 5*μM*] and *U*_*down*_ ∈ [0.25*μM*, 2*μM*] are randomly selected within their value ranges.

In the training stage, the optimization target is the relationship between phasic/tonic responses and the input strength(Fig.2C&2D). Specifically, in a single trace of responses, the tonic response *R*_*tonic*_ is the stable response at the end, and the phasic response *R*_*phasic*_ is the maximum response minus *R*_*tonic*_. To qualitify the performance of a model, we collected 20 traces of responses and calculated *R*_*phasic,i*_, *R*_*toninc,i*_, maximum Ca2+ concentration *U*_*top,i*_ and resting concentration *U*_*down,i*_ of each trace, where i=1,…,20. The features error of the model for optimization is

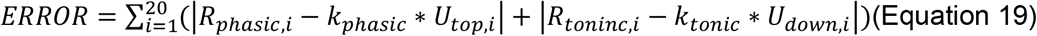

Here *kphasic* = 4(*ves*./*ms*) and *ktonic* = 0.4(*ves*./*ms*) represent relationships between responses and the Ca2+ concentration in experimental recordings.

In the testing stage, the OFF-Response protocols follow the origin paper (Fig.6C in(12)). In this protocol, the membrane potential of the cone begins at −35mV and then goes to its resting potential (−70mV). The resting potential lasts for *Δt* (50*ms*, 150*ms*, 250*ms*, 40*ms*, 700*ms*, 1*s*, 1,5*s*, 2*s*, 3*s*, 4*s*, 5*s* in each). In the end, the potential is elevated to −10mV to trigger a phasic response. This phasic response is named the ‘off-response’ because the membrane potential protocol mimics the moment the light turns dark. We integrated responses at the last stage to represent the strength of the response *R*_*off*_(Fig.S5C). We use the one-exponential function to fit 11 pairs (*Δt, R*_*off*_) and get the recovery time constant *τ*, as

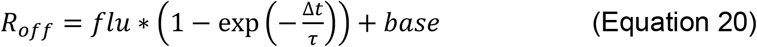

where *flu* indicates the strength of the depression.

### The experiment on the rat rod bipolar cell synapse

All experimental protocols and referenced recordings come from(16) and early related work(50). In the training protocol, the membrane potential begins at −70mV and then goes to *V*_*b*_ for 1000ms. At last, the membrane potential go to −20mV for another 1000ms(Fig.3C). In the N-RAB model, a linear normalization is appended after the RAB module to ensure the maximum amplitude of model responses is the same as the measurement (330pA). Thus, the scaler in the RAB module, N, doesn’t affect the N-RAB model’s performance, so we permanent the N=200 as an RBC-AII synapse has ten active zones(50), and each zone keeps 20 vesicles(27). The integrated transient response *Q*_*trainsent*_ is the sum of the phasic release component, corrected for the previous tonic responses. Each phasic stage is no longer than 500ms.

In the training stage, the error of a model is the sum of absolute differences between the model response’ feature and the corresponding one in recordings, as

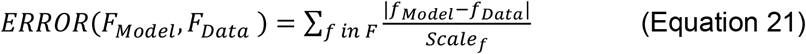

where *F*_*Model*_ and *F*_*Data*_ present the collection of features from simulated and recorded features, *F*_*Model*_ and *F*_*Data*_ indicate the single feature. *Scale*_*f*_ is a factor to unify the unit of features. Specifically, features and corresponding factors are:

1. Integrated transient response *Q*_*trainsent*_ at the 1st stage(Fig.3C(3)). *Q*_*trainsent*_s from model responses are normalized to [0, 1]. A hill function regresses features from recordings, as

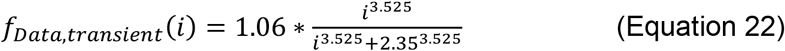

where *i* = 0,1,2,3,4, 5,6 for *V*_*b*_ = *-*55, *-*50, *-*45, *-*40, *-*35, *-*30, *-*25*mV*, respectively. *Scale*_*f*_ = 1 as all features are unit-free.
2. Integrated response before membrane potential goes to −20mV (Fig.S1C). *F*_*Model*_ is the sum of output in 1st stage with a) corrected for the resting response, and b) range normalization. *F*_*Data*_ comes from (Fig,3C, in(16)). Here again, *Scale*_*f*_ = 1.
3. The sum of *Q*_*transient*_ in two stages are close to the same value. *F*_*Model*_ is the ratio between the sum of transient responses at the 1^st^ stage and the 2^nd^ stage. *F*_*Data*_ = 1. Here again, *Scale*_*f*_ = 1.
4. The proportion of transient response in total response, *F*_*Model*_ is the *Qtransient* at the 1st stage in *V*_*b*_ = *-*25*mV*, divided by the total integrated response. *F*_*Data*_ = 1/4.6. *Scale*_*f*_ = 1.
5. The 2^nd^ peak response in the trace *V*_*b*_ = *-*55*mV* is the same as the 1^st^ peak response in the trace *V*_*b*_ = *-*25*mV. fModel* is the maximum response of the trace at *V*_*b*_ = *-*25*mV* divided by the one at *V*_*b*_ = *-*55*mV. fData* = 1. *Scale*_*f*_ = 1.
6. Phasic release time course at *V*_*b*_ = *-*25*mV*. *F*_*Model*_ is the time course of the decay curve at the 1st. *F*_*Data*_ = 5*ms*. *Scale*_*f*_ = 5(*ms*) for normalization.

In the testing stage, the paired-pulse depression protocol comes from Fig.5A (50). In the original experiment, the resting potential is −60mV, and the pulse potential is +90mV. However, we cannot use the +90mV as it is out of the membrane voltage *V*_*m*_ range. In our experiments, we used the −10mv from the Fig.1(50) to trigger transient release. Each pulse lasts 10ms, and the interval between two pulses varies from 50ms to 15s. The recovery response here is the maximum amplitude of the 2nd peak, corrected with the response at the resting potential. The analysis method for recovery time constants is the same as in the salamander cone synapse experiment (Equation 20).

### Compared Models

In the stage of model comparison, we select five models from previous works:

A. The Linear-Nonlinear-Kinetics (**LNK**) Model It is an adaptive model that mimics the vesicle process inside the synapse(49). The linear filter and the nonlinearity are adopted from the LN block’s linear temporal filter L, and the sigmoid *N*_*LNS*_ function (Equation 14). We test all types of kinetics blocks, LNK-BC, LNK-OFFGC, and LNK-ONGC, and find that the LNK-ONGC is the best one (Fig.S2).
B. The Generlized Linear Model (**GLM**) It is an adaptive model whose outputs tune the input strength with a linear feedback filter(83). In this work, the GLM model contains two linear temporal filters. The 1^st^ filter and a nonlinearity process the inputs, and the 2^nd^ temporal filter processes the output of the nonlinearity. The response is added to the inputs for the 1^st^ temporal filter. Notably, this GLM model differs from previous GLM models in the continuous response instead of digital spikes. The linear filter and the nonlinearity are the same with the LNK model.
C. The ribbon synapse model in the original paper(the **Oesch-Origin** model(16)) This model is a single-pool vesicle release model, similar to the RAB module, but with two differences: a) The membrane voltage directly changes vesicles’ release/resupple probability. b) Formulations of release/resupply probabilities, as

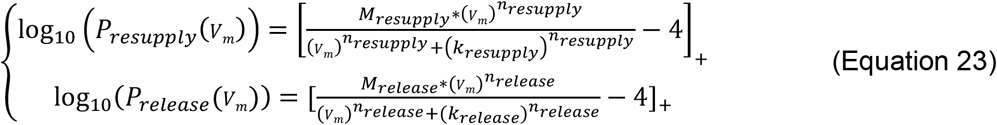

where *P*_*resupply*_ and *P*_*release*_ represents probabilities of resupply and release. *M*_*resupply*_, *n*_*resupply*_, *k*_*resupply*_, *M*_*release*_, *n*_*release*_, *k*_*release*_ are parameters. In this case, parameters are adapted from the original paper without optimization.
D. Same with C), but parameters are free in optimization (the **Oesch-Opt** model(16))
E. A ribbon synapse model with cascade vesicle pools (the **Schroder** model(30)) It is a biophysical model for ribbon synapses with vesicle movement rates among three vesicle pools and an endocytosis rate from released vesicles. It has three vesicle pools (RP, IP, RRP) and an ‘exo’ state to control the response. Transfer rates among pools are

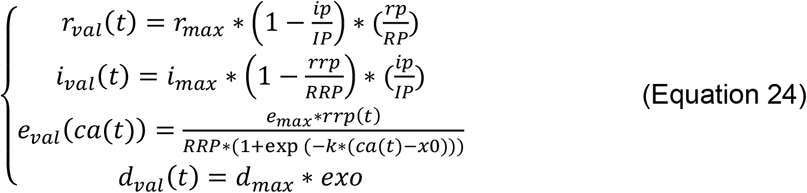

where *r*_*max*_, *i*_*max*_, *e*_*max*_, *d*_*max*_, *k, x*0 are parameters. *RP, IP, RRP* represent the size of vesicle pools, and *rp, ip, rrp* represent the current number of vesicles inside pools. The change of vesicles in three pools and the *exo* state are

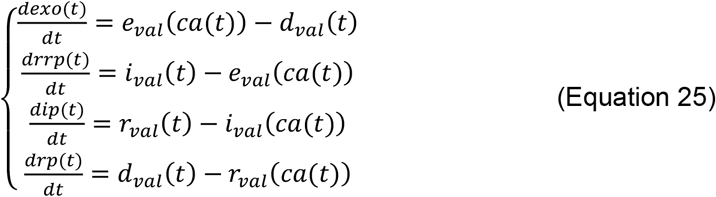

We follow the original paper to fix the value of parameters, where *d*_*max*_ = 0.1 and *RRP* = 35186. Other parameters are free during optimization. Notably, this model receives the Ca2+ concentration instead of the membrane potential. Thus, we used the nonlinearity in the N-RAB model to convert the membrane potential into the Ca2+ concentration (Equation 15). This V-I function is fixed during optimization.

In the training stage, several models have good performance(FigS.3). In contrast, they generate fast recoveries from depression in the testing stage(FigS.4B).

### The experiment on mouse cone bipolar cell synapses

All experimental protocols and referenced recordings come from(35). We used the dataset with local chirp and Strychnine to block the functions of amacrine cells. Besides, we omitted recordings with eight subtypes of BCs for their abnormal, high-noise recordings. The temporal filter at the end of the hLNS model, representing the function of iGluSnFR, is a linear temporal filter *L*(*x*) with a fixed time constant(*τL* = 60*ms*) for all BC-types(28, 58).

In this experiment, the optimization target is the response curve *RData*. We define the error of the model as the sum of the absolute error of the simulated trace *RModel*, as

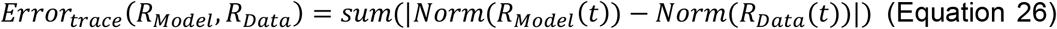

where *Norm* represents the normalization method. It ensures the response rests(the stable response at the beginning) at 0 and its maximum response is 1, as

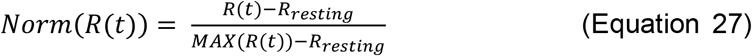

For noise-free simulated traces, *R*_*resting*_ = *R*(*t* = 0). For noisy recordings, the *R*_*resting*_ is the average value of responses before the stimuli start (previous 1000ms).

We define the gain of the RAB module as the change in output (*ΔO*) caused by a small change in the input (*Δ*_*u*_), following(49), as

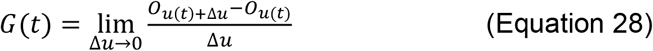

We can derive the instantaneous gain at any moment when *Δ*_*u*_ tends to be 0. Through the definition of output (Equation 3), we get the relationship between gain and parameters, as

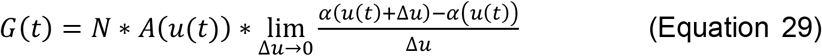

In Equation 30, the third part is the derivation of *α*. Thus, the gain of the RAB block, as Equation 7. In the experimental approach to measure the gain, a small pulse lasts for 10ms with a perturbation *Δ*_*u*_ = 0.01.

### Simulations and optimizations

We applied all models and related models in the Python2/3 programming environment. For K units in LNK models, we applied them in the NEURON simulation platform and its Python API. Each model has 100 (except 1000 for the hLNS model) initial sets of parameters within value ranges and optimized by the ‘Nelder-Mead’ method in *scipy*.*minimize* (84). We select the best one for later analysis. The timestep of simulation is 1ms in all cases.

All models are steady before simulating, except the Schroder model, which cannot analyze the steady state. For the Schroder model, we simulate it 4000ms before the beginning, following the original paper(30). In the simulation of the RAB module, we use an explicit solver instead of numerical procedures. The solver for linear differential Equation 3 is

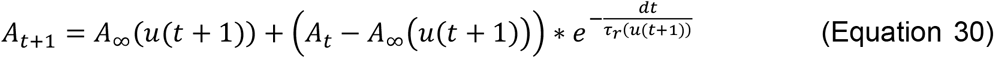

