## Supplemental Tables and Figures for "A computational framework linking synaptic adaptation to circuit behaviors in the early visual system"

Table S1. Variables in the RAB module and analogous biophysical processes.

| variable in the RAB module | Analog to the biophysical process | Experiments with Measurements | Notes |
|---|---|---|---|
| $u(t)$ | The Ca2+ concentration | /, the input. | Several experiments and measurements used voltage clamps to manipulate the membrane potential. |
| $O(u(t))$ | The vesicle release rate of the RRP | /, the output. | Several experiments and measurements recorded the postsynaptic currents. |
| $\alpha(u(t))$ | The release rate constant for a single vesicle | (1, 2) | / |
| $\beta(u(t))$ | The resupply rate constant for a single vesicle | /, unmeasurable. | (3, 4) suggest the calcium signal controls the vesicle resupply process. |
| $O_\infty(u(t))$ | The stable vesicle release rate | (4, 5) | (4) recorded the postsynaptic currents to estimate the presynaptic vesicle release rate. |
| $A_\infty(u(t))$ | The occupancy rate of vesicles sites in the RRP during the stable release | (6, 7) | Those experiments controlled the membrane potential instead of the Ca2+ concentration. |
| $\tau_r(u(t))$ | The time constant of adaptive response trace | (8, 9) etc. | Those experiments only measure values at specific membrane potentials. |
| $N$ | The number of sites in the RRP | (5, 10, 11) etc. | Those experiments and models assumed sites are filled with vesicles at resting membrane potential. |
| $N * A_\infty(u(t))$ | The number of vesicles in the RRP when the synapse reaches stable | 1) EM (5, 12)<br>2) Electrophysical experiments(9) | 1) EM can estimate the value in dark-adapted or light-adapted synapses.<br>2) Electrophysical experiments can estimate the value at resting, assuming that the high membrane potential(-10mV) can deplete all vesicles in the RRP. |

Table S2. Formulations of state transfer rates in all candidates for the RAB module.

| | Linear rate(1) $\alpha(u) = k_{RL} * u$ | Hill rate(2) $\alpha(u) = \frac{m_{RS} * u^{n_{RS}}}{{k_{RS}}^{n_{RS}} + u^{n_{RS}}}$ | Linear stable(5) $O_\infty(u) = k_{OL} * u$ | Power stable(4) $O_\infty(u) = A_{OP} * u^{B_{OP}} + C_{OP}$ |
|---|---|---|---|---|
| Linear $A_\infty(u) = k_{AL} * u + 1$ | $\beta(u) = -k_{RL} * u * \left(\frac{k_{AL} * u + 1}{k_{AL} * u}\right)$ <br> $\tau_r(u) = \frac{-k_{AL} * u}{k_{RL} * u}$ | $\beta(u) = \frac{-m_{RS} * u^{n_{RS}}}{{k_{RS}}^{n_{RS}} + u^{n_{RS}}} * \left(\frac{k_{AL} * u + 1}{k_{AL} * u}\right)$ | $\alpha(u) = \frac{k_{OL} * u}{N * (k_{AL} * u + 1)}$ <br> $\beta(u) = -\frac{k_{OL} * u}{N * k_{AL} * u}$ | $\alpha(u) = \frac{A_{OP} * u^{B_{OP}} + C_{OP}}{N * (k_{AL} * u + 1)}$ <br> $\beta(u) = -\frac{A_{OP} * u^{B_{OP}} + C_{OP}}{N * k_{AL} * u}$ |
| Exponential $A_\infty(u) = \exp(u/k_{AE})$ | $\beta(u) = k_{RL} * u * \left(\frac{\exp(u/k_{AE})}{1 - \exp(u/k_{AE})}\right)$ | $\beta(u) = \frac{m_{RS} * u^{n_{RS}}}{{k_{RS}}^{n_{RS}} + u^{n_{RS}}} * \left(\frac{\exp(u/k_{AE})}{1 - \exp(u/k_{AE})}\right)$ | $\alpha(u) = \frac{k_{OL} * u}{N * \exp\left(\frac{u}{k_{AE}}\right)}$ <br> $\beta(u) = \frac{k_{OL} * u}{N * \left(1 - \exp\left(\frac{u}{k_{AE}}\right)\right)}$ | $\alpha(u) = \frac{A_{OP} * u^{B_{OP}} + C_{OP}}{N * \exp\left(\frac{u}{k_{AE}}\right)}$ <br> $\beta(u) = \frac{A_{OP} * u^{B_{OP}} + C_{OP}}{N * \left(1 - \exp\left(\frac{u}{k_{AE}}\right)\right)}$ |
| Hill $A_\infty(u) = \frac{{k_{AH}}^{n_{AH}}}{{k_{AH}}^{n_{AH}} + u^{n_{AH}}}$ | $\beta(u) = k_{RL} * u * \left(\frac{{k_{AH}}^{n_{AH}}}{u^{n_{AH}}}\right)$ | $\beta(u) = \frac{m_{RS} * u^{n_{RS}}}{{k_{RS}}^{n_{RS}} + u^{n_{RS}}} * \left(\frac{{k_{AH}}^{n_{AH}}}{u^{n_{AH}}}\right)$ | $\alpha(u) = \frac{k_{OL} * u}{N} * \frac{{k_{AH}}^{n_{AH}} + u^{n_{AH}}}{{k_{AH}}^{n_{AH}}}$ <br> $\beta(u) = \frac{k_{OL} * u}{N} * \frac{{k_{AH}}^{n_{AH}} + u^{n_{AH}}}{u^{n_{AH}}}$ | $\alpha(u) = \frac{A_{OP} * u^{B_{OP}} + C_{OP}}{N} * \frac{{k_{AH}}^{n_{AH}} + u^{n_{AH}}}{{k_{AH}}^{n_{AH}}}$ <br> $\beta(u) = \frac{A_{OP} * u^{B_{OP}} + C_{OP}}{N} * \frac{{k_{AH}}^{n_{AH}} + u^{n_{AH}}}{u^{n_{AH}}}$ |
| Sigmoid $A_\infty(u) = \frac{1}{1 + \exp\left(-\frac{u - half_{AS}}{slope_{AS}}\right)}$ | $\beta(u) = k_{RL} * u * \frac{1}{\exp\left(-\frac{u - half_{AS}}{slope_{AS}}\right)}$ | $\beta(u) = \frac{m_{RS} * u^{n_{RS}}}{{k_{RS}}^{n_{RS}} + u^{n_{RS}}} * \left(\frac{1}{\exp\left(-\frac{u - half_{AS}}{slope_{AS}}\right)}\right)$ | $\alpha(u) = \frac{k_{OL} * u}{N} * \left(1 + \exp\left(-\frac{u - half_{AS}}{slope_{AS}}\right)\right)$ <br> $\beta(u) = \frac{k_{OL} * u}{N} * \left(1 + \exp\left(-\frac{u - half_{AS}}{slope_{AS}}\right)\right) * \frac{1}{1 + \exp\left(-\frac{u - half_{AS}}{slope_{AS}}\right)}$ | $\alpha(u) = \frac{A_{OP} * u^{B_{OP}} + C_{OP}}{N} * \left(1 + \exp\left(-\frac{u - half_{AS}}{slope_{AS}}\right)\right)$ <br> $\beta(u) = \frac{A_{OP} * u^{B_{OP}} + C_{OP}}{N} * \left(1 + \exp\left(-\frac{u - half_{AS}}{slope_{AS}}\right)\right) * \frac{1}{1 + \exp\left(-\frac{u - half_{AS}}{slope_{AS}}\right)}$ |

Table S3. Formulations in previous single-pool ribbon synapse models.

| $\beta$ \ $\alpha$ | Constant | Linear | Hill | Quadratic | Sigmoid |
|---|---|---|---|---|---|
| Constant | (10, 13) | (5) | (14) | (15) | (16) |
| Hill | | | (7) | | |

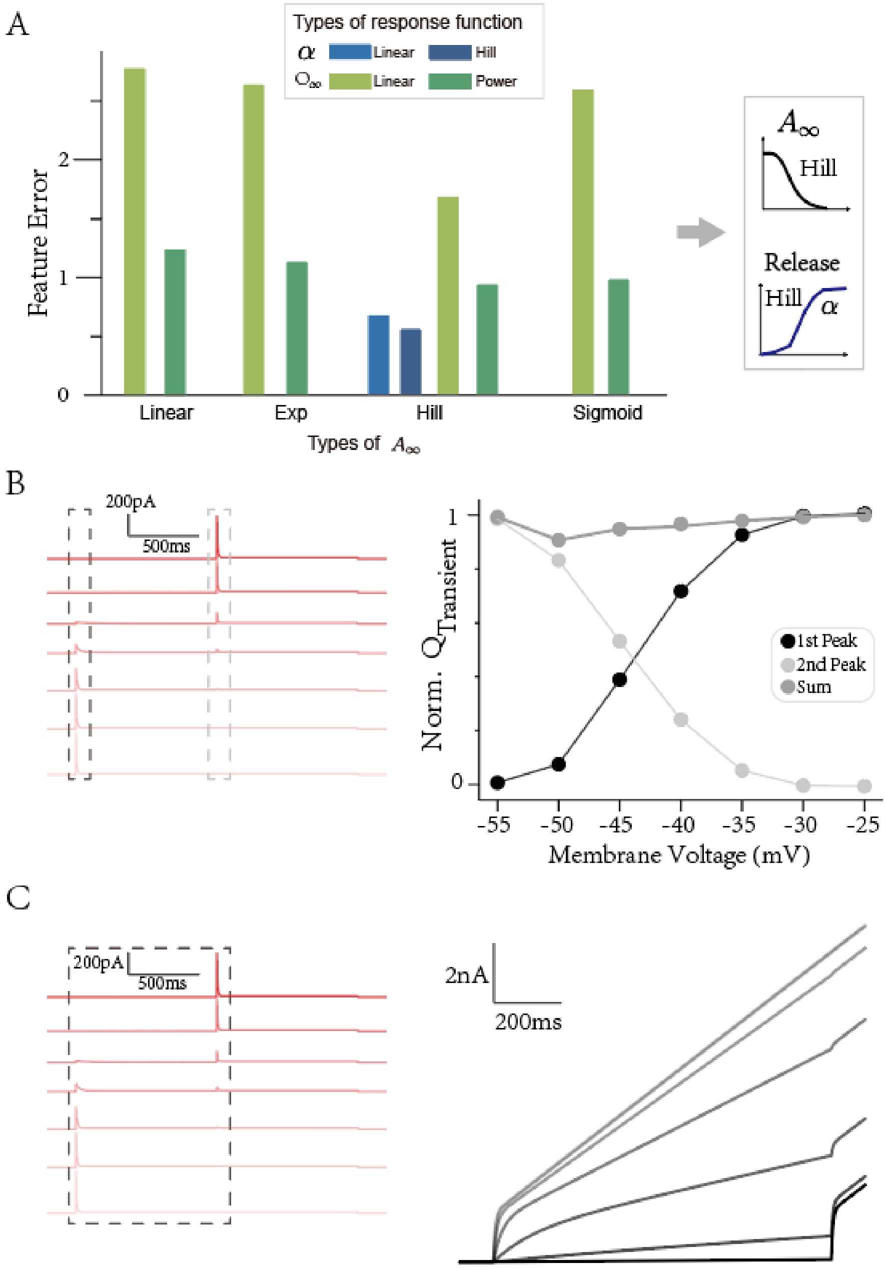


**Figure S1. Parameter Selection in the rat rod bipolar cell (Fig.3) and related features.** (A) The performance of candidates (left) and the best (right). Several candidates don't have suitable parameters for minimal requirements of response traces(see Methods). (B) The feature is that the sum of two transient responses(left) is almost the same among traces(right), mimicking the original experimental figure 3B in(7). (C) The integrated responses until the transient response of the 2nd stage ends, mimicking the original experimental figure 3C in(7).

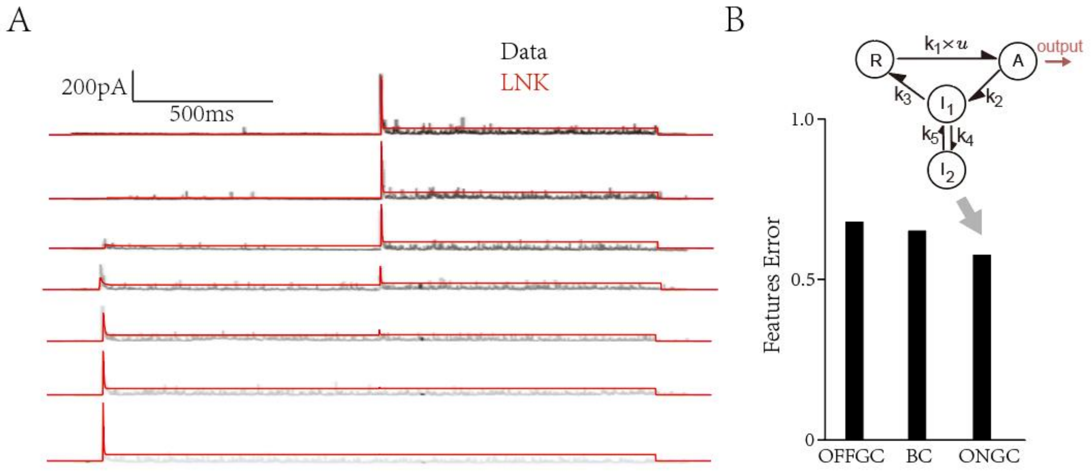


**Figure S2. The performance of the LNK model in the rat rod bipolar cell synapse (Fig.3).** (A) Simulated responses from the optimized LNK-ONGC model. (B) The LNK model with the ONGC-type kinetics block (up) has the best data-fitting performance (down).

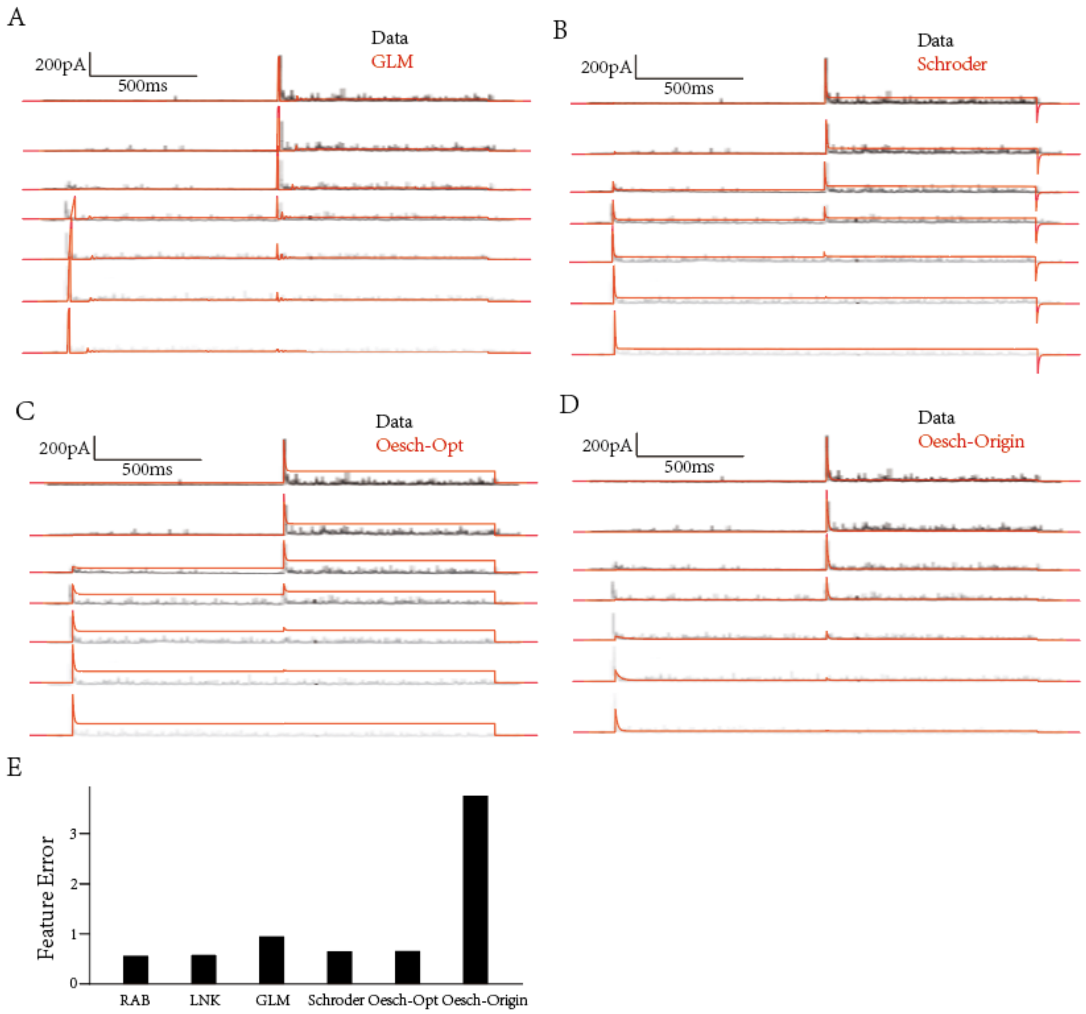


**Figure S3. The performance of other adaptive or ribbon synapse models in the rat rod bipolar cell (Fig.3).** (A) the GLM model, (B) the Schroder ribbon synapse model, (C) the model in the original paper (7) with optimization, and (D) the model in the original paper without tuning parameters. (E) The summary of model performances.

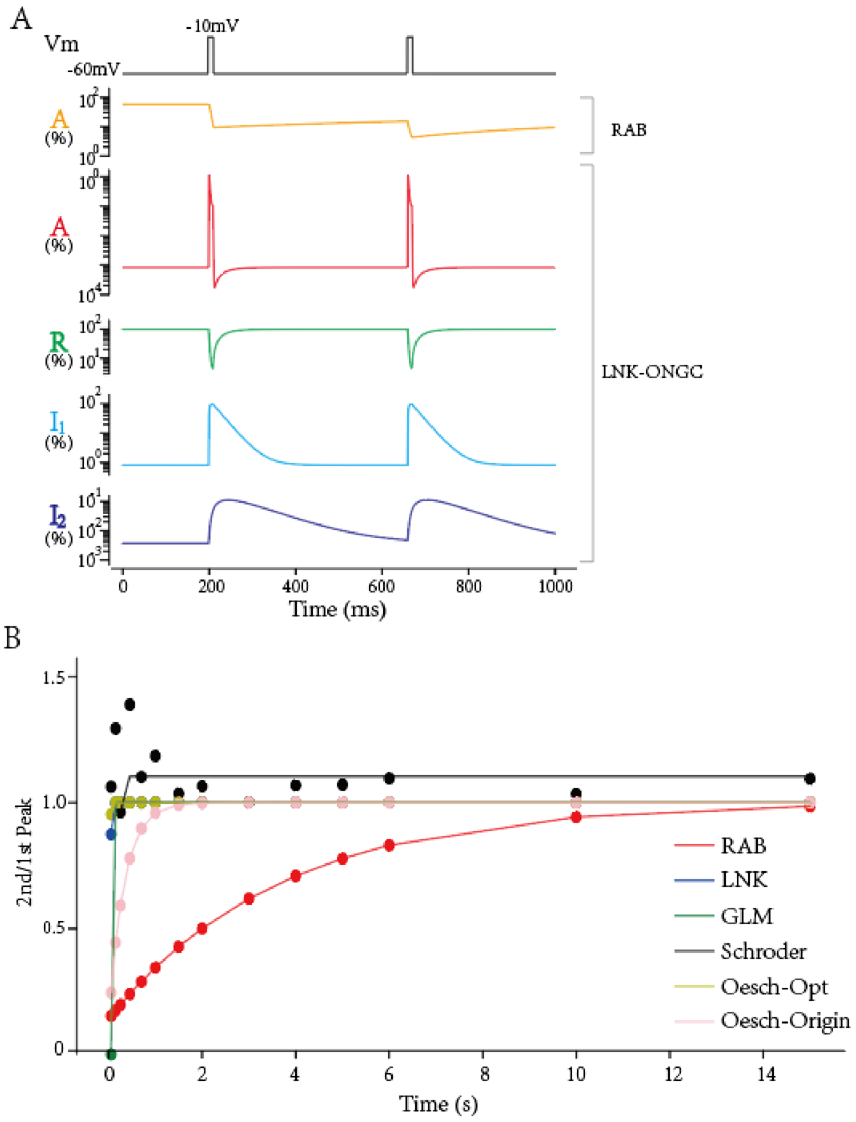


**Figure S4. Inner dynamics at the PPD protocol (Fig.4).** (A) Traces of internal states of the RAB module(up) and the ONGC-type block(down) at the PPD protocol ($\Delta t = 250ms$). (B) Summary of depression recoveries on all computational models in Figure S3.

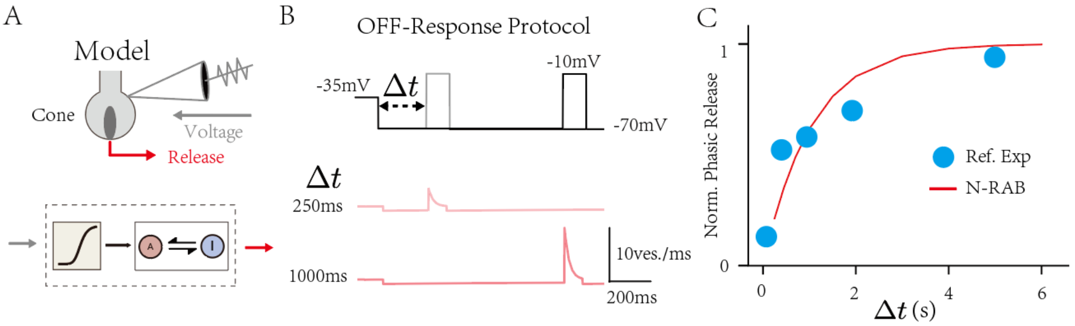


**Figure S5. The adaptation testing for the short-term depression of the salamander cone synapse (Fig.2).** (A) The N-RAB mode of the cone synapse. (B) The OFF-response protocol. (up) It mimics how the synapse adapts to light (-35mV) and produces an 'off-response' (-10mV), adapted from Figure 6C in(5). (down) The responses of the cone synapse model at input trace with $\Delta t = 250ms$ and $\Delta t = 1000ms$. (C) The recovery of off-responses in the model (red) and the experimental recordings (blue, adapted from Figure 6D in(5)).

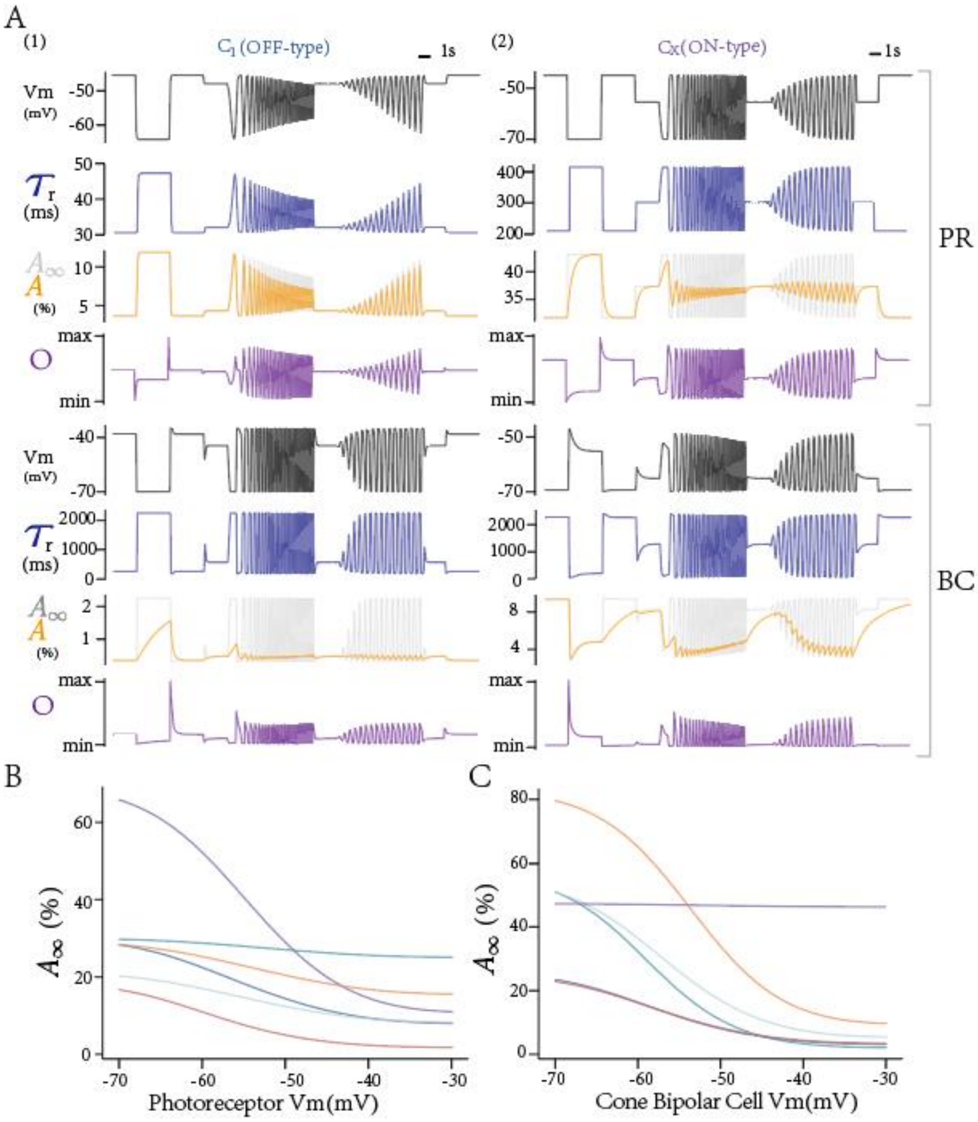


**Figure S6. Inner dynamics in the hLNS model (Fig.5).** (A) Traces of variables in (1) an on-type $C_1$ and (2) an off-type circuit $C_X$. (B and C) The stable states of the active state A in two layers of RAB modules.

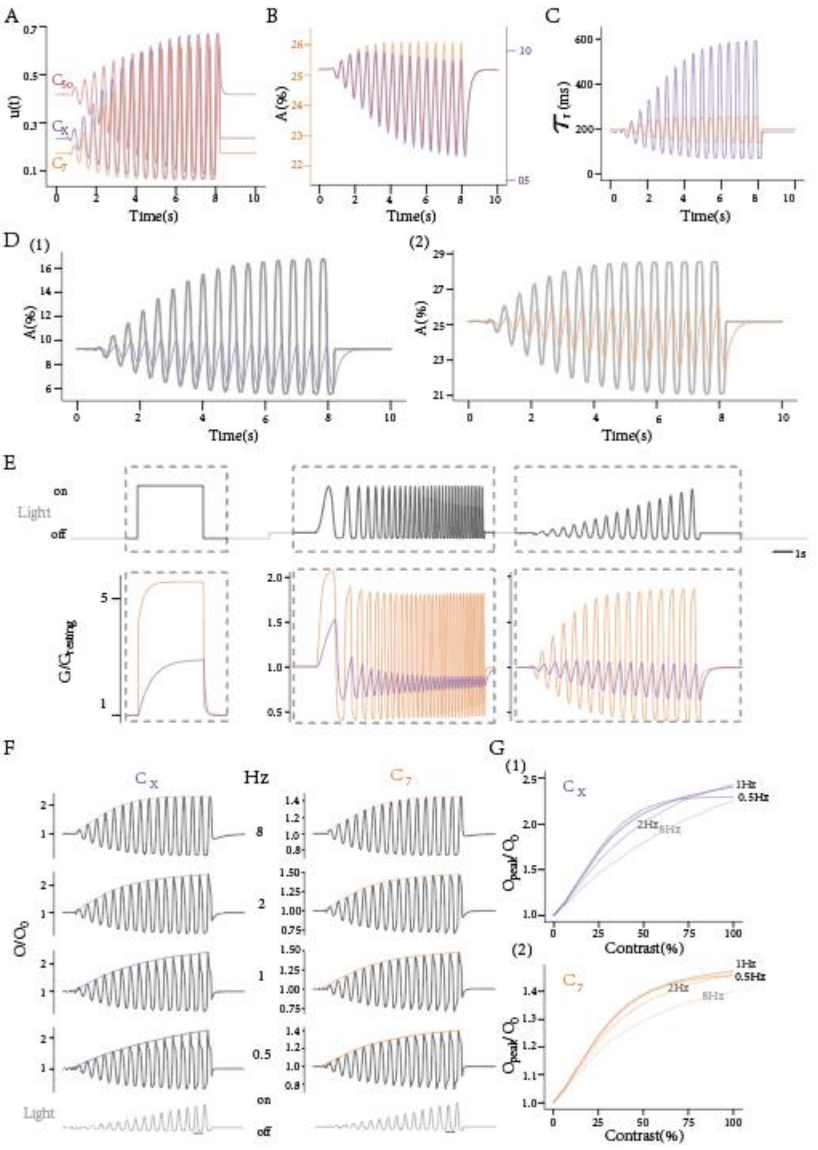


**Figure S7. Gain analysis of the RAB module in the hLNS model (Fig.6).** (A) Inputs for the 1st RAB module in the $C_{5o}$ circuit differ from other on-type ones. (B and C) Traces of stable states and time constants in the 1st RAB module of the $C_X$ and the $C_7$ circuit. (D) Traces of the current state of A in the $C_X$ and the $C_7$ circuit. (E) Gain controls in other experimental stages. (F and G) The output control in the two modules is similar.
